# Wee1 opposes APC/C^Cdh1^ activity to promote S-phase entry

**DOI:** 10.64898/2025.12.01.691493

**Authors:** Luc-Alban Vuillemenot, Lucas G. Morales, Calin-Mihai Dragoi, Betheney R. Pennycook, Edward Harris, Liuyang C. Lin, William A. Weston, Jade-Ellen Brown, Emma Graham, Ivan Andrew, Georgia Roumelioti, Alex Montoya, Laurence Game, George R. Young, Pavel Shliaha, Bela Novak, Alexis R. Barr

## Abstract

Wee1 phosphorylates and inhibits CDK activity to inhibit mitotic entry and establish a G2 DNA damage checkpoint. Consequently, Wee1 inhibitors are in clinical trials, developed to be synthetically lethal in *TP53* mutant tumours that become reliant on a Wee1-mediated DNA damage checkpoint. However, Wee1 inhibitors have efficacy in *TP53* wild-type tumours and many trials have been terminated due to high levels of toxic side-effects, suggesting that Wee1 has unknown functions. Here, we show that Wee1 promotes cell cycle re-entry from quiescence (G0) by opposing the activity of the E3 ubiquitin ligase, APC/C^Cdh^^1^. Wee1 phosphorylates Cdh1 (*FZR1*) at key residues that mediate the interaction between Cdh1 and APC/C. Cells with loss-of-function of Wee1 during G0/G1 have delayed S-phase entry, an impaired G1/S transition, abnormal S-phase accumulation of the CDK inhibitor p21 and enter a p21-dependent G2 arrest. Reduced expression of APC/C^Cdh^^1^ or p21 renders cells more sensitive to acute Wee1 inhibition and both pathways are downregulated in acquired Wee1 inhibitor resistance. Our study reveals a new cell cycle control mechanism that has implications for how Wee1 inhibitors should be used in the clinic.

## Summary

Cyclin-dependent kinases (CDKs) control passage through the eukaryotic cell cycle. Wee1 is a tyrosine kinase that inhibits CDK1 activity during G2 by phosphorylating Tyr15 in the CDK1 active site ^1–3^. Late in G2, Cdc25 phosphatases dephosphorylate Tyr15, promoting CDK1 activation and mitotic entry ^4–9^. This mechanism forms a G2 DNA damage checkpoint, which becomes vital in *TP53* mutant cancer cells ^10–12^. Therefore, Wee1 inhibitors were developed to be synthetically lethal in *TP53* mutant cancers and are currently being trialled across a range of cancer types ^13–17^. However, many trials are being suspended due to toxic side-effects ^17^, the origins of which are not fully understood. Wee1 also phosphorylates and inhibits CDK2 that regulates G1 progression and S-phase entry. CDK2^T14A;Y15F^ knock-in mutant mice and human colorectal cancer cells enter S-phase faster when returning to the cycle from quiescence ^18,19^. However, injection of IMR90 human fibroblasts with a plasmid expressing Wee1 kinase had no effect on S-phase entry timing ^20^ and there is only indirect evidence that Wee1 impacts CDK2 activity in cells ^21^. Therefore, the role of Wee1 in regulating S-phase entry is incompletely understood and we sought to clarify the function(s) of Wee1 in early cell cycle control. Here, we reveal a new role for Wee1 in suppressing APC/C^Cdh^^1^ activity during G1 to promote timely S-phase entry in cells returning to proliferation from quiescence and identify Cdh1 as a new Wee1 substrate important for cell cycle control. Our findings shed light on the pleiotropic impact of Wee1 inhibitors (Wee1i) in cells.

## Results

### Wee1 is required for efficient S-phase entry

To understand the potential roles of Wee1 in regulating CDK2 activity during G1 and S-phase entry, we constructed an influence diagram describing the known interactions of Wee1 with the G1/S regulatory network (Figure 1A). Wee1 has two inputs into this network through inhibition of CyclinE/CDK2 and CyclinA/CDK2 ^22,23^. We converted this influence diagram to a series of differential ODEs and parametrised the model based on known dynamics of the G1/S regulatory network (Figure 1B, ^24,25^). To identify the role of Wee1 in this network, we simulated the depletion of Wee1 levels to 50% of baseline. Subsequently, we observed that CDK2 is activated earlier than in the presence of 100% Wee1 activity and that S-phase entry is accelerated.

**Figure 1.**
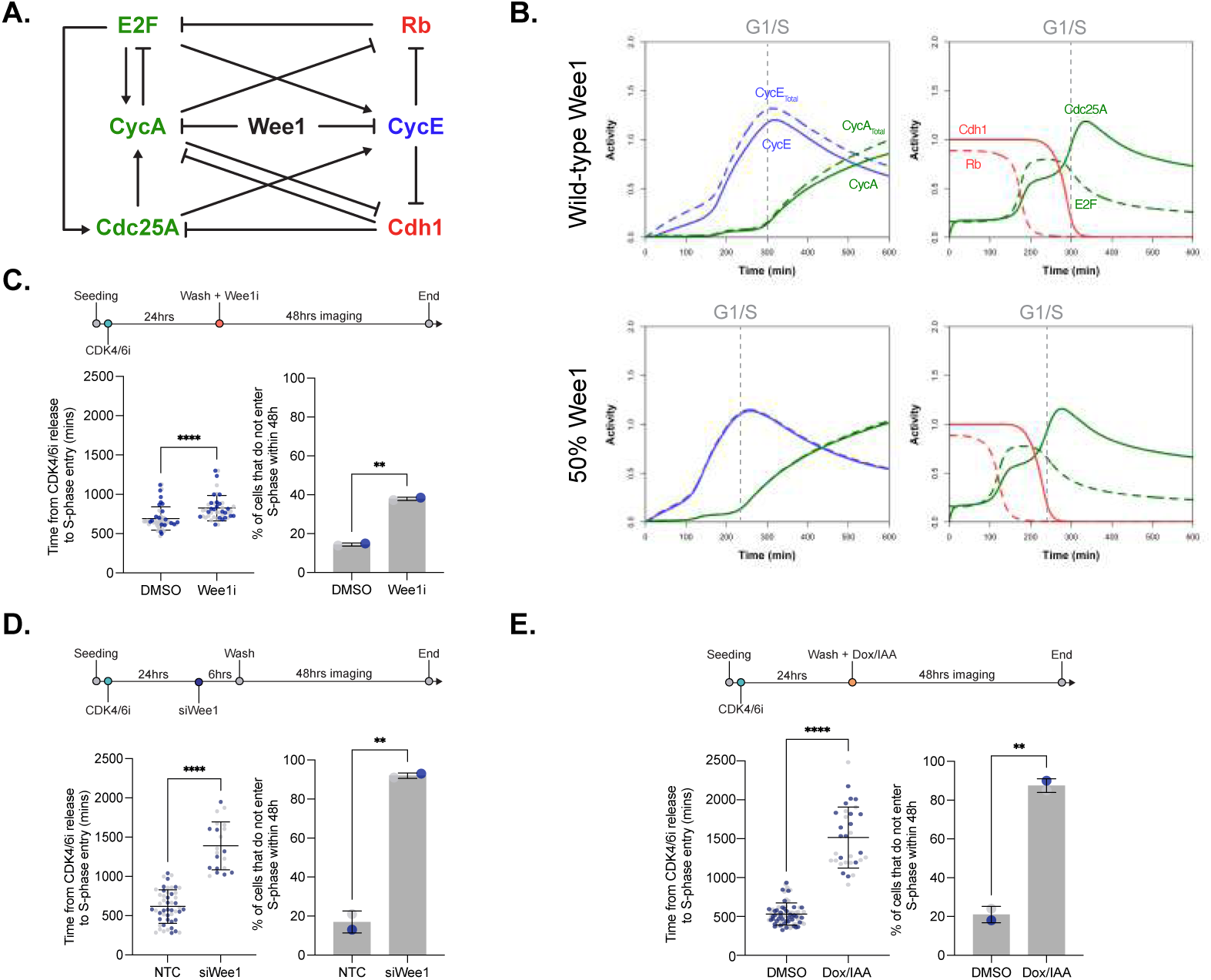
Wee1 is required for efficient S-phase entry **A.** Influence diagram of the G1/S regulatory network. **B.** Time course simulation of the ODE model demonstrating how reducing Wee1 activity to 50% of wild-type levels predicts a faster increase in CDK2 activity and a shorter G1 phase. CycE_Total_ is Tyr15- phosphorylated (inactive) plus Tyr15-unphosphorylated (active) CDK2 combined (blue dashed curve). CycE is only Tyr15-unphosphorylated CDK2 (active, blue solid curve). The same logic applies to CycA in green. The G1/S transition is marked with a grey dashed line, determined by the time of APC/C^Cdh1^ switch off (red solid curve) and increase in CycA/CDK2. **C.** Graphs show length of G1 phase (left) and fraction of cells that do not enter S-phase (right) after Wee1i treatment upon release from CDK4/6i- mediated G0/G1 arrest. Experimental timeline shown above the graphs. **D.** Graphs show length of G1 phase (left) and fraction of cells that do not enter S-phase (right) after Wee1 depletion by siRNA 6h before release from CDK4/6i-mediated G0/G1 arrest. Experimental timeline shown above the graphs. **E.** Graphs show length of G1 phase (left) and fraction of cells that do not enter S-phase (right) after Wee1 degradation by addition of doxycycline (Dox) and IAA upon release from CDK4/6i- mediated G0/G1 arrest. Experimental timeline shown above the graphs. In C-E, all experiments are n=2, mean +/- stdev are plotted with black lines, individual data points are shown in blue or grey, coloured by experimental repeat. Unpaired student’s t-test for significance. ****p<0.0001; **p<0.01.

To experimentally test whether we observe accelerated S-phase entry in cells when Wee1 activity is reduced, we used the Wee1 inhibitor (Wee1i), AZD1775, currently in clinical trials. We selected a dose of 1 μM, which is sufficiently high to inhibit Wee1, but minimises off-target effects (Figure S1A-D). To investigate the role of Wee1 in S-phase entry while avoiding confounding effects from Wee1 perturbations during the preceding G2, we synchronised cells in early G0/G1. For this, we treated hTert-RPE1 mRuby-PCNA cells^26^ with the CDK4/6 inhibitor (CDK4/6i), Palbociclib (Figure S1E; ^27^). Subsequently, we washed out CDK4/6i, released the cells into G1 in the presence or absence of Wee1i and quantified G1 length by timelapse imaging of mRuby-labelled PCNA. Contrary to our model predictions and current understanding of Wee1 functions, Wee1 inhibition increased G1 length and the fraction of cells that did not enter S-phase during the 48h imaging period (Figure 1C). To determine that this was not an artefact of CDK4/6i treatment, we also arrested cells in G0 by contact inhibition followed by serum starvation (Figure S1F; ^28^). Then, we replated cells at low density, in the presence or absence of Wee1i, released them back into the cell cycle and quantified G1 length by timelapse imaging. Again, we observed increased G1 length and an increased fraction of cells not entering S-phase after Wee1 inhibition (Figure S1G).

Since kinase inhibitors can have off-target effects, we depleted Wee1 from CDK4/6i arrested cells using siRNA (Figure S1H), then released them back into the cell cycle and quantified G1 length by timelapse imaging. Wee1 depletion had an even more severe impact than Wee1i, where very few cells entered S-phase over the 48h imaging period, and those that did, had an increased G1 length (Figure 1D). To be certain that the delay in entering S-phase was due to loss of Wee1 function, we also generated a cell line with an inducible Wee1-GFP-degron (Figure S2A-E) to remove Wee1 in a highly-specific manner. We validated that this cell line had the expected nuclear localisation of Wee1-GFP (Figure S2B), that it grew with normal kinetics (Figure S2D) and that Wee1 could be degraded within 1 hour upon the addition of doxycycline and IAA (Figure S2E). Degradation of Wee1 upon release from a CDK4/6i-mediated G0/G1 arrest again led to most cells not entering S-phase and, of those that did, G1 length was significantly longer in the absence of Wee1 (Figure 1E).

Together, these data reveal that, contrary to our predictions and current understanding of Wee1 functions, Wee1 inhibition does not accelerate S-phase entry. Instead, we find that Wee1 is required for timely S-phase entry in cells returning to proliferation from quiescence.

### Wee1 inhibition leads to uncoupling of CDK2 activity and S-phase entry

In addition to predicting a shorter G1 length upon Wee1 inhibition, our model (Figure 1A,B) predicted an earlier and more rapid increase in CDK2 activity. Therefore, we wanted to determine if and how CDK2 activity changes during G1 when Wee1 is inhibited. We made use of a fluorescent live-cell reporter (CDK2L-GFP) that measures CyclinE/CDK2 activity during G1 ^24^ to quantify CDK2 activity in real-time (Figure 2A). hTert-RPE1 mRuby-PCNA CDK2L-GFP cells were arrested in G0/G1 with CDK4/6i and released into S-phase in the presence or absence of Wee1i, while CDK2 activity was tracked by timelapse imaging. We opted to use Wee1i to perturb Wee1 since this is a clinically relevant drug and a higher fraction of cells enter S-phase, compared to Wee1 siRNA or degron approaches, allowing us to make statistically meaningful comparisons. When Wee1 is inhibited, CDK2 activity starts to increase earlier during G1 and reaches a higher activity than in DMSO-treated cells (Figure 2B; Supplementary Movie 1). Cells treated with Wee1i therefore enter S-phase at a higher level of CDK2 activity. These results were recapitulated using a similar, shorter version of the CDK2 sensor labelled with mCherry (DHB-mCherry; Figure S3A,B; ^29^).

**Figure 2.**
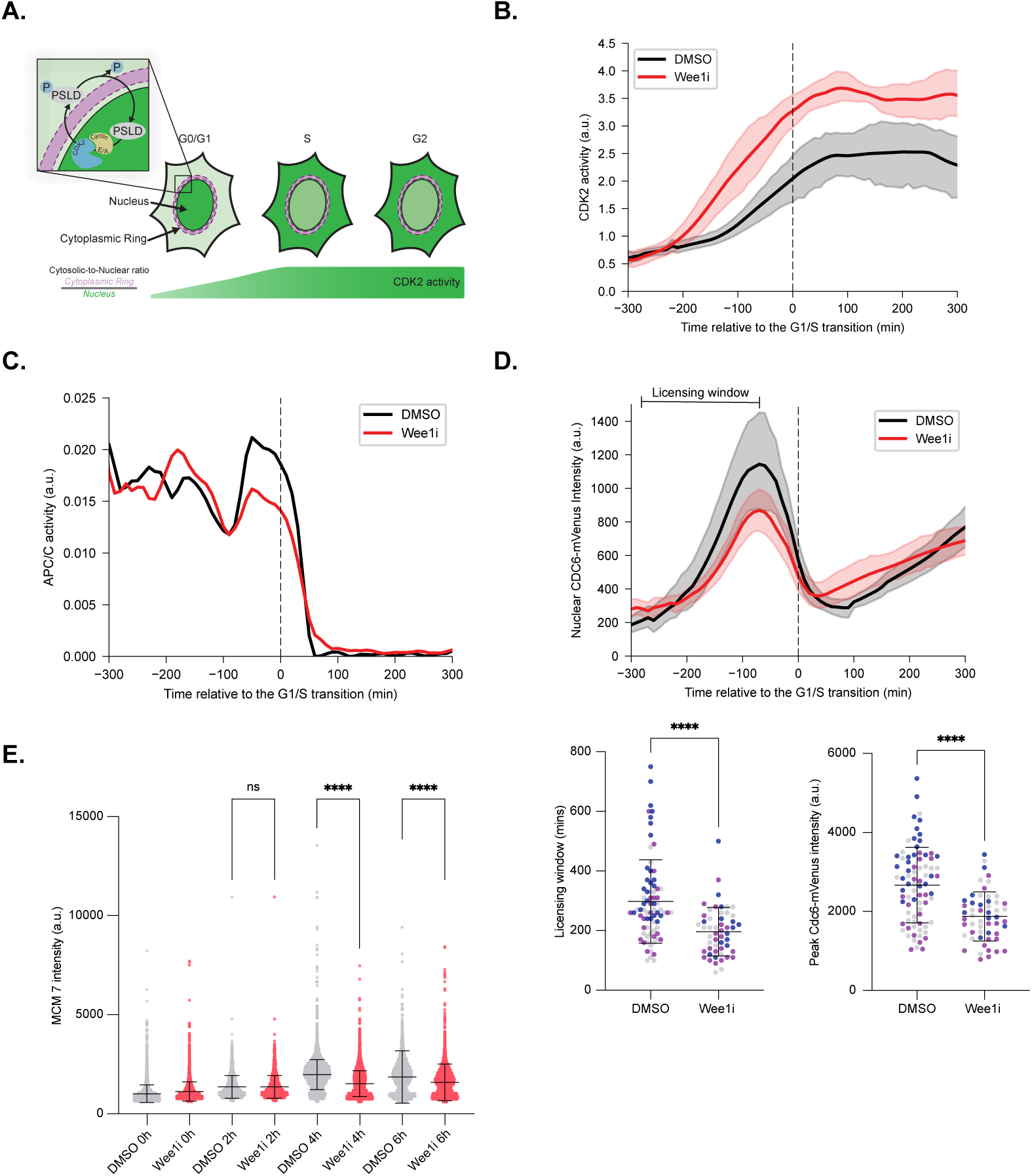
Wee1 inhibition leads to uncoupling of CDK2 activity and S-phase entry. **A.** Cartoon showing how the CDK2L-GFP CDK2 activity sensor works. **B.** Graph shows CDK2 activity plotted relative to the G1/S transition for cells released from G0/G1 arrest in the presence of DMSO (control, black line) or Wee1i (red line). Mean and 95% confidence interval around the mean over time is shown. **C.** Graph shows timing and rate of APC/C^Cdh1^ inactivation based on a mCherry-Geminin-based readout, plotted relative to the G1/S transition for cells released from G0/G1 arrest in the presence of DMSO (control, black line) or Wee1i (red line). Mean over time is shown. **D.** Upper graph shows expression of Cdc6-mVenus plotted relative to the G1/S transition for cells released from G0/G1 arrest in the presence of DMSO (control, black line) or Wee1i (red line). Mean and 95% confidence interval around the mean over time is shown. The licensing window is denoted, as per^28^. Lower graphs show measurements of the length of the licensing window defined by Cdc6 expression (left graph) and peak Cdc6-mVenus intensity (right graph). Mean +/- stdev are plotted with black lines, individual data points are coloured by experimental repeat. Unpaired student’s t-test for significance. ****p<0.0001. **E.** Graph shows quantification of chromatin-bound MCM7 protein at time intervals after release from CDK4/6i-mediated G0/G1 arrest in the presence of DMSO (control, grey points) or Wee1i (red points). Mean +/- stdev are shown. One-way ANOVA followed by Šídák’s multiple comparisons test for significance. ****p<0.0001; ns = not significant.

Two key events determine S-phase entry, CDK2 activity increase and the inhibition of the APC/C^Cdh^^1^ E3 ubiquitin ligase activity (Figure 1B; ^30^). CDK2 can phosphorylate Cdh1 and inhibit APC/C^Cdh^^1^ activity ^31–33^, raising the question of when APC/C^Cdh^^1^ is switched off in Wee1i-treated cells that have high CDK2 activity but delayed S-phase entry. To answer this question, we made use of an hTert-RPE1 cell line expressing mCherry-Geminin^1–110^ under the control of an exogenous promoter ^34^ as a live-cell reporter of APC/C^Cdh^^1^ activity ^32^. Using this reporter, we identified that in cells released from a CDK4/6i-mediated G0/G1 arrest, APC/C^Cdh^^1^ activity is switched off just after the G1/S transition in Wee1i-treated cells, with indistinguishable timing and rate from that observed in DMSO-treated cells (Figure 2C, S3C). We observed a very similar result when using CyclinA2 as a readout of APC/C^Cdh^^1^ activity, since CyclinA2, like Geminin, is an APC/C^Cdh^^1^ substrate (Figure S3D,E; ^35^). These data suggest that, upon G1 inhibition, CDK2 activity increases more rapidly and to a higher level, but the timing and rate of APC/C^Cdh^^1^ inactivation relative to the G1/S transition remain similar to those in DMSO-treated cells. Consequently, APC/C^Cdh^^1^ inactivation is more important in establishing the timing of S-phase entry than a threshold of CDK2 activity.

Low CDK2 activity during G1 provides a period for DNA replication origin licensing^36,37^. Therefore, we wondered if the early increase in CDK2 activity in G1 in Wee1i-treated cells leads to a decrease in DNA replication origin licensing. Cdc6 is an essential replication origin licensing factor, required to form pre-replicative complexes at replication origins ^38^. Thus, we made use of an hTert-RPE1 cell line expressing exogenous mTurquoise2-PCNA, DHB-mCherry and Cdc6-mVenus ^28^ to track the dynamics of Cdc6 protein, relative to the G1/S transition. In cells released from CDK4/6i G0/G1 arrest in the presence of the Wee1i, we observed that the licensing window (time from the rise to the peak in Cdc6-mVenus intensity ^28^) was reduced, compared to DMSO-treated cells (Figure 2D). Considering these data, we also quantified replication origin licensing during G1 by quantifying the levels of chromatin-bound MCM protein. Consistent with our Cdc6-mVenus timelapse data, we observed a decrease in chromatin-bound MCM7 protein in Wee1i-treated G1 cells as compared to DMSO-treated cells (Figure 2E).

In summary, our results demonstrate that Wee1 acts to inhibit CDK2 activity during G1 and that an early and rapid increase in CDK2 activity after Wee1 inhibition can lead to reduced DNA replication origin licensing but not inhibition of APC/C^Cdh^^1^ activity. Increased CDK2 activity is insufficient to accelerate the transition of Wee1i-treated G1 cells into S-phase, suggesting that Wee1 plays other, unknown, roles during G1.

### S-phase entry is abnormal in Wee1 inhibited cells

Since progression through G1 is abnormal in Wee1i-treated cells, with a prolonged period of combined high CDK2 and high APC/C^Cdh^^1^ activity, we examined if Wee1i- treated cells entered S-phase normally. We observed that mRuby-PCNA localization at the very start of S-phase was abnormal in Wee1i-treated cells with low PCNA intensity and PCNA foci less well-defined (Figure 3A). We quantified the standard deviation of mRuby-PCNA nuclear intensity which showed a clear reduction at the G1/S transition in Wee1i-treated cells compared to DMSO-treated cells (Figure 3B). This decrease remained throughout S-phase, indicating altered S-phase progression in the presence of the Wee1i, consistent with previous reports ^39–41^.

**Figure 3.**
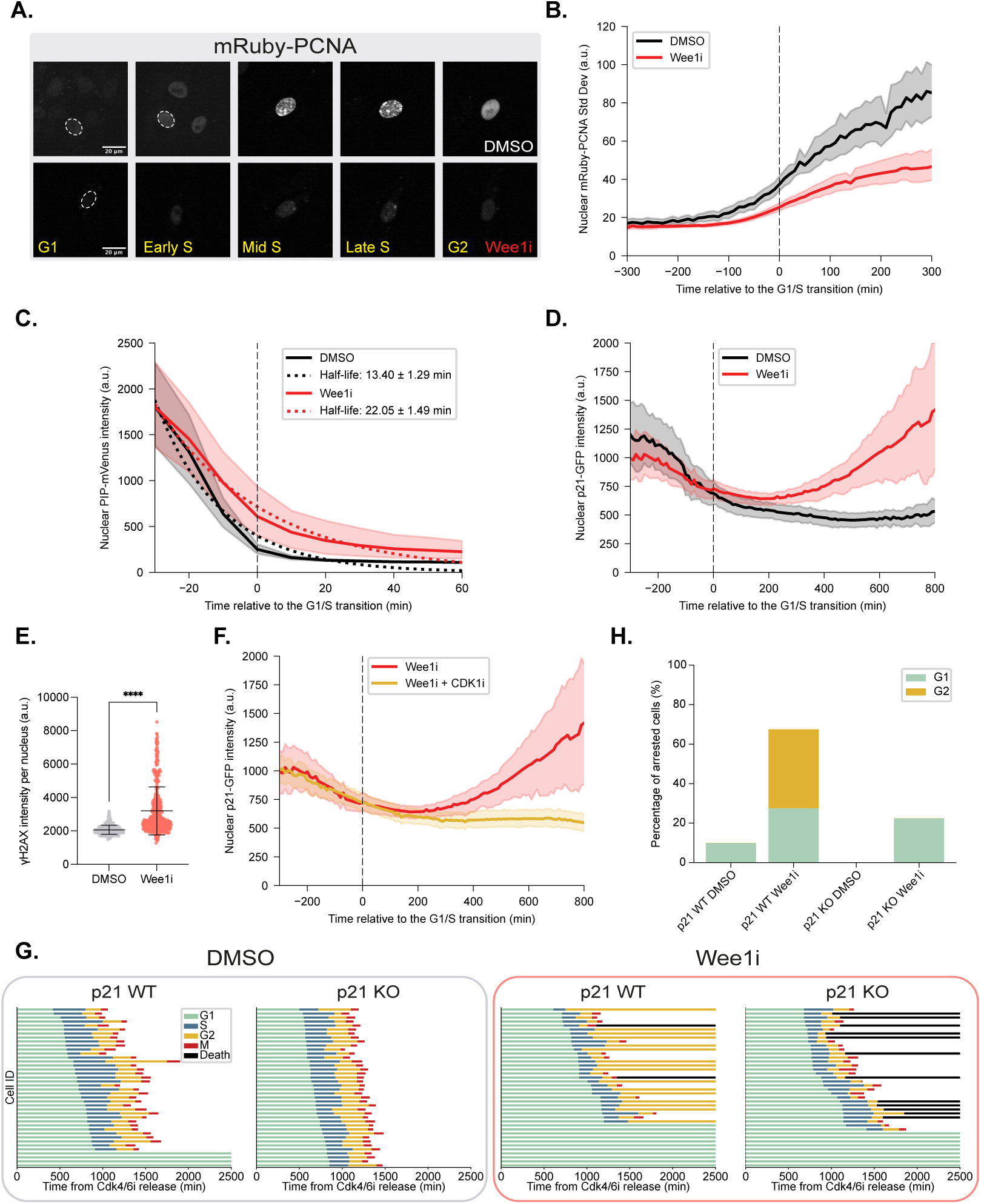
S-phase entry is abnormal in Wee1 inhibited cells **A.** Images showing differences in mRuby-PCNA localisation between DMSO (upper panels) and Wee1i-treated (lower panels) cells. **B.** Quantification of the standard deviation (stdev) of mRuby-PCNA intensity over time, plotted relative to the G1/S transition for cells released from G0/G1 arrest in the presence of DMSO (control, black line) or Wee1i (red line). Mean and 95% confidence interval around the mean over time is shown. **C.** Quantification of PIP-mVenus intensity over time, plotted relative to the G1/S transition for cells released from G0/G1 arrest in the presence of DMSO (control, black line) or Wee1i (red line). Mean and 95% confidence interval around the mean over time is shown. One-phase exponential decay is denoted by the dotted lines and half-life for each condition is shown. **D.** Quantification of p21-GFP nuclear intensity over time, plotted relative to the G1/S transition for cells released from G0/G1 arrest in the presence of DMSO (control, black line) or Wee1i (red line). Mean and 95% confidence interval around the mean over time is shown. **E.** Graph shows quantification of nuclear gH2Ax intensity in EdU-positive cells released from G0/G1 arrest for 48h in the presence of DMSO or Wee1i. Student’s t-test for significance. ****p<0.0001. **F.** Quantification of p21-GFP nuclear intensity over time, plotted relative to the G1/S transition for cells released from G0/G1 arrest in the presence of Wee1i (red line) or combined Wee1i and CDK1i (yellow line). Mean and 95% confidence interval around the mean over time is shown. **G.** Graphs show cell cycle phase timings for p21 wild-type (WT) and p21 knockout (KO) cells treated with DMSO (left graphs) or Wee1i (right graphs) after release from CDK4/6i. **H.** Graph shows fraction of cells arresting in G1 or G2 phase in p21WT and p21KO cells treated with DMSO or Wee1i.

PCNA recruitment to chromatin in S-phase brings substrates, including the DNA licensing factor Cdt1 and the CDK inhibitor protein p21, in contact with the CRL4^Cdt^^2^ E3 ubiquitin ligase. This occurs through substrate-binding to PCNA via a PCNA interaction protein (PIP) motif ^42^, which leads to rapid substrate degradation and S-phase progression. We assayed whether the change in PCNA recruitment we observed in Wee1i-treated cells impacts PIP-containing substrate degradation by using an hTert-RPE1 cell line expressing the PIP-FUCCI reporter ^34^ and timelapse imaging. This construct contains the PIP motif of Cdt1 fused to an mVenus fluorophore, allowing us to track the dynamics of its degradation at the G1/S transition. In Wee1i-treated cells, the degradation of PIP-mVenus was slower compared to DMSO-treated cells (Figure 3C), suggesting that impaired recruitment of PCNA to chromatin at the start of S-phase impacts the rate of degradation of PIP-containing substrates.

p21 contains a PIP motif and is normally rapidly degraded at the start of S-phase ^43–45^. If p21 is not degraded during S-phase, it can cause severe disruption to DNA replication and the accumulation of DNA damage ^46^. Considering our findings in the PIP-FUCCI cell line, we wanted to see if p21 degradation was impaired after Wee1 inhibition. We used an hTert-RPE1 mRuby-PCNA cell line expressing endogenously labelled p21-GFP ^25^ to track p21 dynamics in real-time. We observed that p21-GFP protein accumulates abnormally during S-phase (Figure 3D; Figure S4A,B; Supplementary Movie 2). This suggests ongoing impaired p21 degradation in S-phase and potentially, increased p21 synthesis. Transcription of p21 is induced by p53, downstream of DNA damage ^10,47^. We observed that cells treated with Wee1i had increased levels of the DNA damage marker gammaH2AX (Figure 3E, Figure S4C), consistent with known impacts of Wee1 inhibition during S-phase ^41,48,49^. This could drive increased p21 protein synthesis. However, any p21 protein synthesized during S-phase should be degraded by the CRL4^Cdt^^2^ complex. In addition to impaired degradation of p21 in S-phase by reduced PCNA recruitment to chromatin, we hypothesized that Wee1 inhibition in S-phase would lead to increased CDK1 activity and the premature inactivation of the CRL4^Cdt^^2^ E3 ligase, which is CDK1-dependent^50^. Therefore, we tested if treating cells with a CDK1 inhibitor would restore CRL4^Cdt^^2^ activity during S-phase and prevent accumulation of p21. Combining Wee1 and CDK1 inhibition prevented the accumulation of p21-GFP in S-phase cells (Figure 3F; Figure S4D,E).

High levels of p21 protein can cause cell cycle arrest in G2. We tracked the cell cycle fate of hTert-RPE1 mRuby-PCNA cells released from a CDK4/6i G0/G1 arrest and treated with Wee1i. We observed an increase in the fraction of cells arresting in G2 in Wee1i-treated cells compared to DMSO-treated cells and determined that the arrest in G2 was p21-dependent (Figure 3G, H). Increased G1 arrest after Wee1i treatment was not p21-dependent (Figure 3G, H). While we call this a G2 arrest, it has been shown that Wee1i treated cells do not completely replicate their DNA during S-phase, therefore more accurately, this is a “G2-like” arrest ^49^. We confirmed that this was the case by examining the Hoechst sum intensity profiles where we observed a large accumulation of cells between 2n and 4n ploidy specifically in Wee1i-treated cells (Figure S4F,G).

Taken together, our data show that Wee1 inhibition leads to an abnormal G1/S transition that is exacerbated by the detrimental effects of Wee1 on DNA replication.

### Wee1 suppresses APC/C^Cdh^^1^ activity to promote S-phase entry

We wanted to determine what was preventing Wee1 inhibited G1 cells with high CDK2 activity from entering S-phase prematurely. Since APC/C^Cdh^^1^ appears to be inactivated with normal rate and timing around the G1/S transition in Wee1i treated cells, we wondered if Wee1 inhibition could be causing increased, or prolonged, APC/C^Cdh^^1^ activity during G1 that could lead to a delay in S-phase entry. This hypothesis is consistent with a previous report showing that Wee1 physically interacts with, and can inhibit the ubiquitin ligase activity of, APC/C^Cdh^^1^ ^151^.

First, we updated our ODE model to test the impact of Wee1 inhibiting APC/C^Cdh^^1^ on the G1/S regulatory network (Figure 4A). We conceived of three potential mechanisms for this interaction. First, we assumed that Wee1 inactivates Cdh1 through an identical mechanism to that of CDK2 (“CDK2-like model”), i.e. Wee1 phosphorylates the same sites in Cdh1 as CDK2. Thus, Cdh1 hyper-phosphorylation and inactivation would be determined by a simple (weighted) sum of CyclinE/CDK2, CyclinA/CDK2 and Wee1 activities (Figure S5A). This model predicts that Wee1 downregulation to 50% of wild-type activity advances CyclinE/CDK2 activation, but also that CyclinE must accumulate to a much higher level to compensate for Wee1 depletion, leading to delayed Cdh1 inactivation and G1/S entry (Figure S5B). Second, we hypothesised that Wee1 does not inactivate Cdh1 directly but rather increases the affinity of CyclinE/CDK2 for APC/C^Cdh^^1^ (“threshold model”) which has otherwise been shown to be negligible *in vitro* ^31,52^. Again CyclinE/CDK2 activation is advanced. One distinct feature of this model is that APC/C^Cdh^^1^ inactivation dynamics display a two-phase profile upon Wee1 inhibition. First, Cdh1 is phosphorylated gradually by the accumulating CyclinE/CDK2 and second, the remaining Cdh1 is rapidly inactivated once CyclinA starts to accumulate (Figure S5E,F). Third, we hypothesised that Wee1 inactivates Cdh1 independently of CDK2, by phosphorylating separate amino acid residues in Cdh1 (“independent model”). Notably, in this instance, partial Wee1 inactivation results in delayed G1/S entry and greater APC/C^Cdh1^ activity in G1 (Figure S5I,J). Nevertheless, APC/C^Cdh1^ is otherwise inactivated in a sharp, stepwise manner at G1/S under wild-type and 50% Wee1 activity conditions alike.

**Figure 4.**
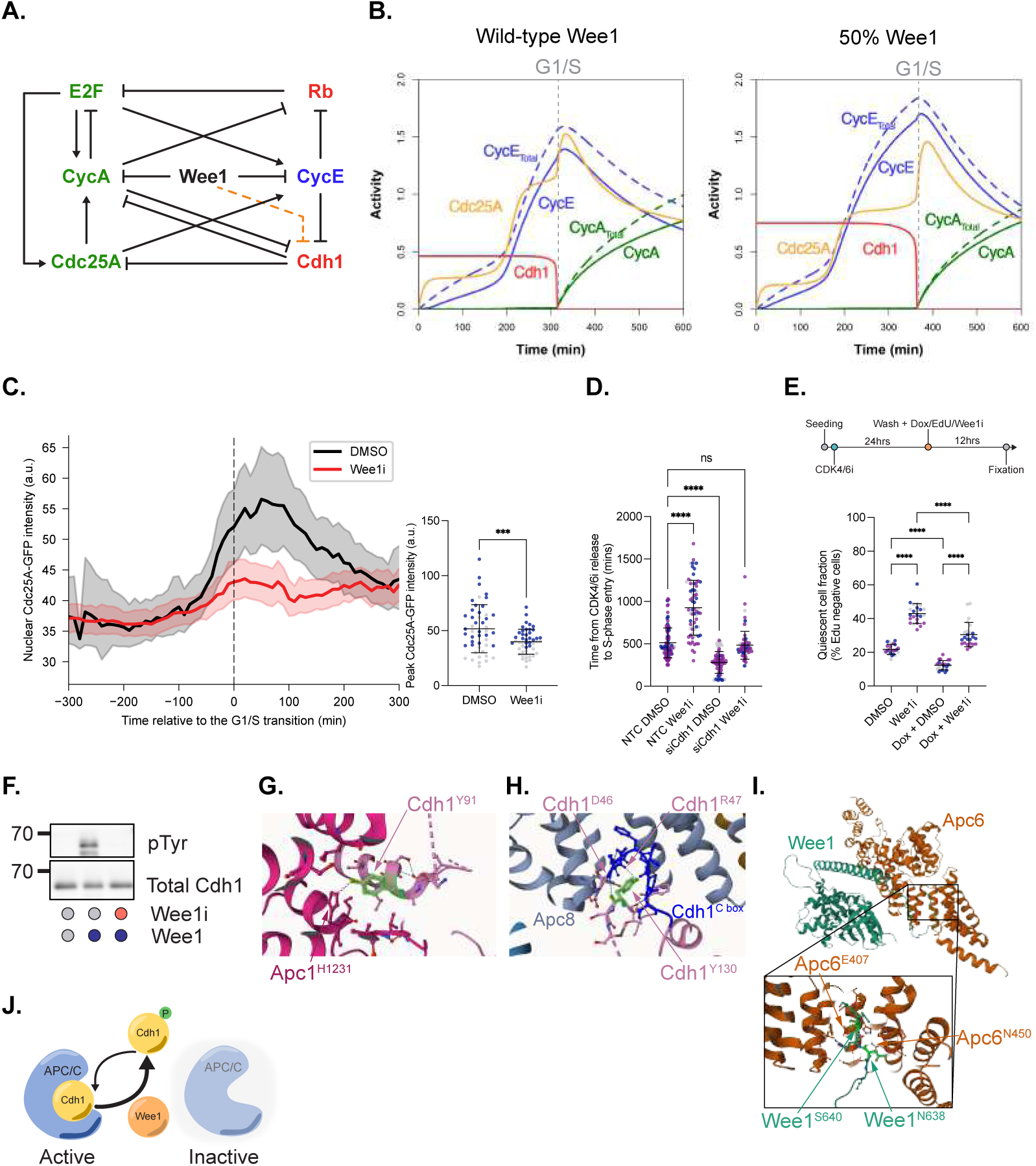
Wee1 suppresses APC/C^Cdh1^ activity to promote S-phase entry **A.** Updated influence diagram of the G1/S regulatory network including the addition of Wee1 inhibiting APC/C^Cdh1^. **B.** Time course simulation from the updated “independent” ODE model demonstrating how reducing Wee1 activity to 50% of wild type levels now predicts a faster increase in CDK2 activity but a longer G1 phase. Also note that during G1, APC/C^Cdh1^ activity is higher under Wee1 inhibition and Cdc25A levels are lower. The G1/S transition is marked with a grey dashed line, determined by the time of APC/C^Cdh1^ switch off and increase in CycA/CDK2. **C.** Quantification of Cdc25A-GFP nuclear intensity over time, plotted relative to the G1/S transition for cells released from G0/G1 arrest in the presence of DMSO (control, black line) or Wee1i (red line). Mean and 95% confidence interval around the mean over time is shown. To the right is the quantification of peak Cdc25A-GFP intensity. Mean +/- stdev are shown in black. Individual data points are coloured by experimental repeat. Unpaired student’s t-test for significance. ***p<0.001 **D.** Graph shows length of G1 phase in cells released from G0/G1 arrest under the conditions labelled. Mean +/- stdev are shown in black. Individual data points are coloured by experimental repeat. One-way ANOVA followed by Tukey’s multiple comparisons test for significance. ****p<0.0001; ns is not significant. **E.** Graph shows fraction of cells arrested in G0/G1 (EdU negative) in cells released from G0/G1 arrest under the conditions labelled. Emi1 refers to doxycycline (Dox)-induced Emi1 overexpression. Cells were released 12 hrs prior to fixation. Mean +/- stdev are shown in black. Individual data points are coloured by experimental repeat. One-way ANOVA followed by Tukey’s multiple comparisons test for significance. ****p<0.0001. Experimental timeline shown above the graph. **F.** Westen blot showing *in vitro* kinase assay for Wee1 and Cdh1. A phospho-tyrosine antibody (pTyr) was used to probe for tyrosine phosphorylation of Cdh1 by Wee1. **G.** APC/C^Cdh1^ structure (PDB9GAW) highlighting the hydrogen bond between Tyr91 in Cdh1 and His1231 in Apc1. **H.** APC/C^Cdh1^ structure (PDB9GAW) highlighting the intramolecular hydrogen bond between Tyr130 and both Asp46 and Arg47 in Cdh1. The Cdh1 C- motif that Asp46 and Arg47 are part of is highlighted in blue to show the interaction with Apc8. **I.** Predicted interaction between Wee1 (Asn290-Ser642) and Apc6 (Met1- Ala532). The zoomed in region shows the interaction between Glu407 and Asn450 in Apc6 with Ser640 and Asn638 in the C-terminus of Wee1. **J.** Cartoon demonstrating how Wee1 phosphorylation of Cdh1 would lead to loss of Cdh1 binding to APC/C and APC/C^Cdh1^ inactivation.

All three models reproduced the G0 arrest upon complete Wee1 depletion (Figure S5D,H,L), similar to that seen after Wee1 siRNA or degradation (Figure 1D,E). However, the three models differ markedly in the G1 APC/C^Cdh1^ patterns of activity emerging during G1 and at the G1/S from partial (50%) Wee1 inhibition. Thus, each set of assumptions makes experimentally verifiable predictions. The dynamics of APC/C^Cdh1^ inhibition at the G1/S in the threshold model (Figure S5F) are inconsistent with what we observed in cells after Wee1 inhibition (Figure 2C, Figure S3E). This leaves the CDK2-like and independent mechanisms of Cdh1 inhibition by Wee1. These two models differ principally in the change in APC/C^Cdh1^ activity during G1 after Wee1 inhibition (Figure S5A,B,I,J). In the CDK2-like model, APC/C^Cdh1^ activity remains unchanged after Wee1 inhibition, though it is prolonged, whereas in the independent model, APC/C^Cdh1^ activity increases after Wee1 inhibition. To differentiate between the two possibilities, we wanted to quantify the level of APC/C^Cdh1^ activity during G1 in DMSO versus Wee1i-treated cells. We used timelapse imaging to quantify the level of APC/C^Cdh1^ substrates that accumulate in G1 as a real-time readout of APC/C^Cdh1^ activity. Two substrates of APC/C^Cdh1^ emerge during G1 – Cdc6 ^53^ and Cdc25A ^54^. Quantifying the peak level of Cdc6-mVenus expression from our previous analysis revealed that Cdc6-mVenus levels are decreased in Wee1i-treated G1 cells, as compared to DMSO-treated cells (Figure 2D). To analyse Cdc25A protein expression in real-time, we generated an hTert-RPE1 mRuby-PCNA cell line where we introduced a GFP fluorophore into the C-terminus of the *CDC25A* gene (Figure S6A). We achieved several heterozygously tagged clones that had a slightly decreased proliferation rate, mainly due to an increase in G1 length (Figures S6B-E). Cdc25A- GFP localised correctly to the nucleus (Figure S6F). Because of this decreased proliferation, we did also try to tag the N-terminus of Cdc25A but were unsuccessful. Therefore, we continued with the Cdc25A-GFP C-terminally tagged cell line. In cells released from CDK4/6i G0/G1 arrest, Cdc25A-GFP showed accumulation around the G1/S transition, as previously described (Figure 4C; Supplementary Movie 3 ^55^). We also confirmed that Cdc25A-GFP is a reliable readout of APC/C^Cdh1^ activity in G1 by depleting Cdh1 by siRNA and quantifying Cdc25A-GFP levels during G1 (Figure S6G). After Wee1i treatment upon release, Cdc25A-GFP levels were much reduced, compared to DMSO-treated cells, around the G1/S transition (Figure 4C). Our results showing decreased Cdc6 and Cdc25A protein after Wee1i during G1 are consistent with increased APC/C^Cdh1^ activity after Wee1 inhibition and the independent model of Wee1 inhibition of Cdh1 (Figure 4B, S5I-L).

To test whether increased APC/C^Cdh1^ activity could be causing a longer G1 phase in Wee1i-treated cells, we used siRNA to deplete Cdh1. Depletion of Cdh1 leads to a short G1 length, as previously reported ^56^ (Figure 4D, Figure S7A). Partial depletion of Cdh1 and simultaneous inhibition of Wee1 restored a normal G1 length, supporting the hypothesis that in Wee1i-treated cells, APC/C^Cdh1^ is causing a prolonged G1 phase (Figure 4D). We also used a second approach to inhibit Cdh1. Emi1 is an APC/C^Cdh1^ inhibitor ^57^. Using an hTert-RPE1 cell line expressing a doxycycline-inducible Emi1^58^, we could see a robust decrease in Cdh1 expression and a decrease in the fraction of cells arresting in G0/G1 in the presence of the Wee1i (Figure 4E, Figures S7B). Of note, we did not observe increased Cdh1 protein expression after Wee1i treatment (Figure S7C), nor did we observe any change on Cdh1 stability in G0-arrested cells treated with Wee1i (Figure S7D,E), suggesting that Wee1 does not regulate Cdh1 protein stability.

Finally, we wanted to see if Wee1 could phosphorylate Cdh1 directly. Using *in vitro* kinase reactions with purified Wee1 and Cdh1 proteins, we observed that Wee1 can phosphorylate tyrosine(s) in Cdh1 (Figure 4F, S7F). Mass spectrometry analysis identified Tyr91 and Tyr130 of Cdh1 as being phosphorylated by Wee1. No phosphorylation at these sites was detected when Wee1 was omitted or when Wee1 was added together with 0.5 µM Wee1i. To increase confidence in these findings, we re-analysed the samples using a targeted LC-MS approach, focusing acquisition on the identified phosphorylated peptides. This provided exceptionally high-quality fragmentation data, allowing unambiguous localisation of the phosphorylation sites and representing the first direct demonstration of Tyr91 and Tyr130 phosphorylation of Cdh1 by Wee1 (Figure S7G,H,). These sites are conserved across species, particularly Tyr130 (Figure S8), and both phosphorylation events have been detected in cells ^59,60^.

Scrutinising the structure of human APC/C^Cdh1^ ^61^, both sites are part of a Cdh1 N-terminal extension that interacts extensively with APC/C. Tyr91 sits at an interface between Cdh1 and Apc1 and forms a hydrogen-bond with His1231 in Apc1 (Figure 4G). Tyr130 forms intramolecular hydrogen-bonds with Asp46 and Arg47 and appears to stabilise the C-box motif in Cdh1 ^62^ (DRFIP) known to be crucial to maintain interaction with Apc8b ^63^ (Figure 4H). Recently improved protein-protein interaction predictions ^64^ predict a high-confidence interaction between the extended Wee1 C-terminus and Apc6 (Figure 4I), providing a way for Wee1 to dock onto the complex and phosphorylate Cdh1.

Our data and modelling propose a new mechanism of G1/S control whereby Wee1 phosphorylates and inhibits APC/C^Cdh1^ (Figure 4J), independently of CDK2-mediated inhibition of Cdh1, to control G1 length and promote robust S-phase entry.

### p21 and APC/C^Cdh1^ expression levels regulate sensitivity to Wee1 inhibitors

Given that Wee1 inhibition leads to increased APC/C^Cdh1^ activity and p21 expression, we wanted to investigate what implications this would have for Wee1i treatment in the clinic.

To see if Wee1i had similar phenotypes in cancer cells as in non-transformed hTert- RPE1 cells, we repeated experiments in *TP53* wild-type KU1919 bladder cancer and MCF7 breast cancer cell lines. We observed that these cells also underwent a significant G1 and G2 arrest after treatment with the Wee1i, similar to phenotypes observed in hTert-RPE1 cells, and G1 length increased in Wee1i-treated cells, although not significantly so (Figure 5A,B, S9A-F). G2 arrest was p21-dependent and in the absence of p21 we observed increased cell death during G2 in Wee1i-treated cells, compared to DMSO-treated cells, in all cell lines tested (Figure 3G, 5A, C, S9A). These data suggest that cells lacking p21 would be more sensitive to Wee1 inhibition. The gene encoding p21, *CDKN1A*, is rarely mutated in cancer, apart from bladder cancer, where between 10-15% of patients have a predicted loss-of-function mutation in p21^65^. These mutations occur early and are present even in normal bladders^66^. Therefore, we wanted to determine if Wee1i treatment could be useful in eradicating pre-tumorigenic *CDKN1A* mutant lesions to prevent a subset of bladder cancers. We generated p21 knockout cell lines in the non-transformed bladder urothelial cell line, HBLAK and found that these cells were indeed more sensitive to Wee1i than their wild-type counterparts (Figure 5D, S9G).

**Figure 5.**
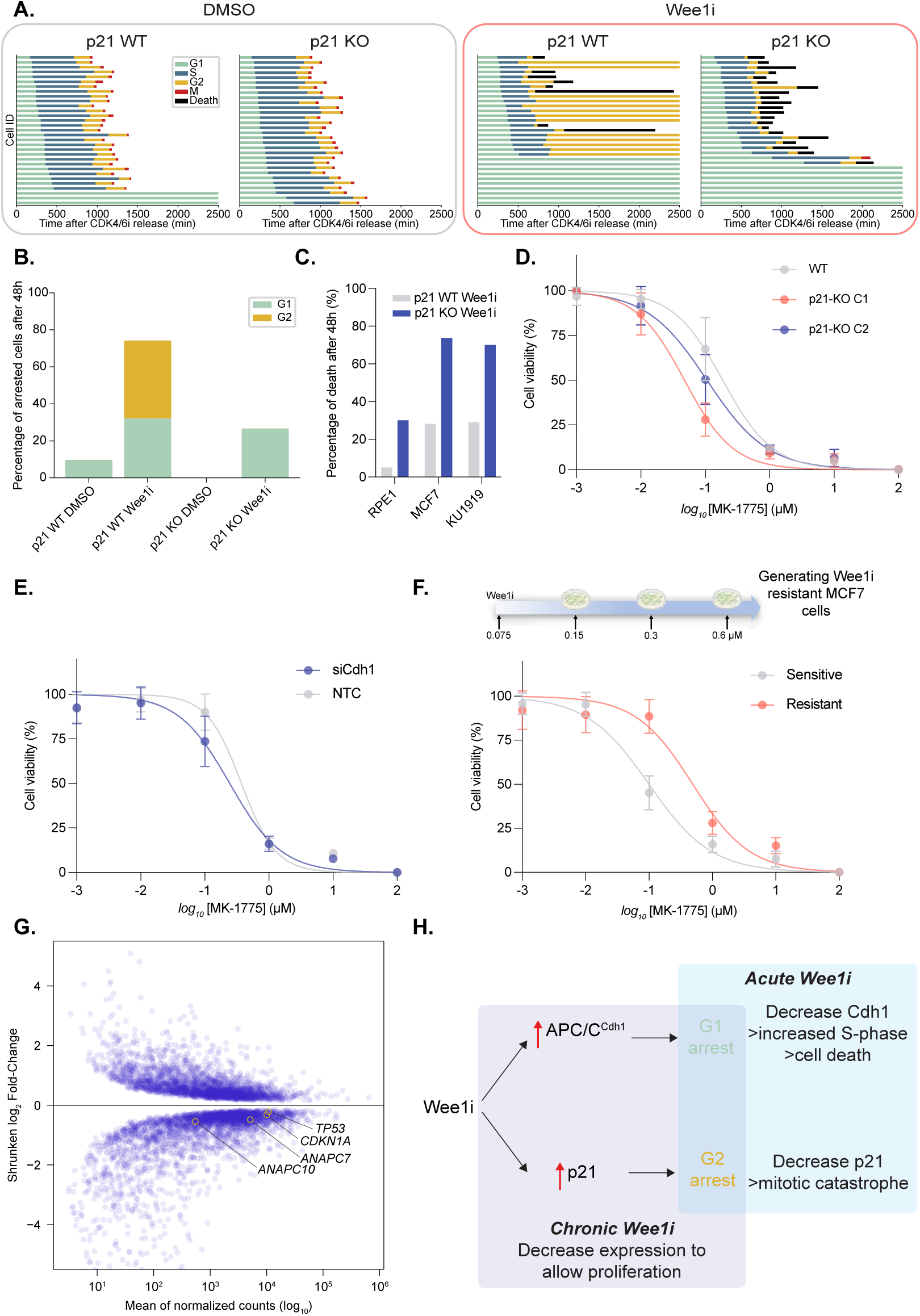
p21 and APC/C^Cdh1^ expression levels regulate sensitivity to Wee1 inhibitors **A.** Graphs show cell cycle phase timings for p21 wild-type (WT) and p21 knockout (KO) KU1919 bladder cancer cells treated with DMSO (left) or Wee1i (right), after release from CDK4/6i. **B.** Graph shows fraction of cells arresting in G1 or G2 phase in p21WT and p21KO cells treated with DMSO or Wee1i from cells plotted in (A). **C.** Graph shows percentage of cells dying, quantified by timelapse imaging, in p21WT and p21KO hTert-RPE1, MCF7 and KU1919 cells treated with DMSO or Wee1i. **D.** Dose response curves show sensitivity of p21WT and p21KO (two clones, c1 and c2) non-transformed HBLAK bladder cell lines to Wee1i treatment (extra sum-of-squares F test, p21WT vs p21KO: p<0.0001). **E.** Dose response curves show sensitivity of cells treated with NTC (grey) or Cdh1- (blue) targeting siRNA, 6 hr prior to treatment with Wee1i (extra sum-of-squares F test; p<0.0001). **F.** Dose response curves show sensitivity of parental (sensitive) and Wee1i resistant MCF7 cells to Wee1i treatment (extra sum-of-squares F test, parental vs Wee1i resistant ; p<0.0001). **G.** Graph shows genes significantly up- and down-regulated (blue) in Wee1i-resistant cells compared to parental (sensitive) cells from RNAseq analysis. *TP53*, *CDKN1A*, *ANAPC7* and *ANAPC10* are highlighted. **H.** Schematic showing the relationship between acute and chronic Wee1 inhibition (Wee1i) with Cdh1 activity and p21 levels.

We reasoned that depletion of Cdh1 may also sensitise cells to Wee1i by allowing those cells to enter S-phase and incur Wee1i-mediated DNA damage. Indeed, siRNA-mediated depletion of Cdh1 increased sensitivity of hTert-RPE1 cells to Wee1i (Figure 5E).

To identify changes that could drive acquired resistance to Wee1i, we generated an MCF7 breast cancer cell line that was resistant to Wee1i (Figure 5F). We profiled these cells by RNAseq and identified pathways that were significantly altered in Wee1i- resistant cells (Figure S9H-J; Supplementary table 1). Wee1i-resistant cells upregulate nucleotide biosynthetic pathways, consistent with previous work showing that nucleoside supplementation reduces double-strand breaks caused by increased replication initiation in Wee1i-treated cells ^39^. Wee1i-resistant cells downregulated *TP53* and *CDKN1A* expression, as well as components of APC/C^Cdh1^ – *ANAPC7* and *ANAPC10* (encoding Apc7 and Apc10, respectively; Figure 5G). These observations are consistent with our mechanistic findings reported here, in that cells would need to downregulate the p53-p21 and APC/C^Cdh1^ pathways to enable continued proliferation in the presence of Wee1i. However, loss of these proteins would lead to increased sensitivity to acute Wee1i treatment, as we see for p21 and Cdh1 (Figure 5C,E,H). Analysing DepMap for genes that synergise with Wee1 loss, also identified several APC/C^Cdh1^ components (Figure S9K).

In conclusion, our data suggest that *CDKN1A* mutation status or APC/C^Cdh1^ subunit expression could lead to improved patient selection and potentially reduced toxicities in Wee1i clinical trials.

## Discussion

Cell cycle-targeting in cancer treatment has regained much enthusiasm in recent years after the success of CDK4/6 inhibitors in breast cancer treatment^67^. While Wee1i were designed to be synthetically lethal with *TP53* mutations for cancer therapy, they are being trialled in both *TP53* wild-type and mutant disease. The significant side-effects associated with Wee1i treatment reported in clinical trials, including myelosuppression, gastrointestinal issues and fatigue, have been attributed to potential off-target effects on Plk1. However, an alternative explanation is that Wee1 has unappreciated functions beyond regulating the G2/M transition. Here, we report a role for Wee1 in regulating cell cycle re-entry from quiescence. This previously unappreciated function of Wee1 could account for some of the impacts of Wee1i observed in healthy tissues.

Previous work had shown that CDK2AF mutant cells, where CDK2 cannot undergo inhibitory phosphorylation by Wee1, have a shorter G1, due to increased CyclinE/CDK2 activity ^68,69^. Therefore, we were initially surprised that Wee1 inhibition, depletion or degradation caused a G1 delay and cells to arrest in G0/G1. However, our data are consistent with these observations since we also observe increased CyclinE/CDK2 activity after Wee1 inhibition. What our experiments reveal is that because Wee1 also inhibits APC/C^Cdh1^ activity, the increase in APC/C^Cdh1^ activity prevents Wee1i-treated cells with high CDK2 activity from entering S-phase prematurely.

Our findings lead to an apparent conundrum in that Wee1 inhibits CDK2 to prevent S-phase entry yet also inhibits APC/C^Cdh1^ to promote S-phase entry. What could be the purpose of these seemingly opposing functions of Wee1 during G1? We argue that Wee1 overall promotes S-phase entry. By inhibiting APC/C^Cdh1^ activity, Wee1 permits Cdc25A expression which reverses Wee1 inhibitory phosphorylation of CDK2. Wee1 may be required for an earlier, initial inhibition of APC/C^Cdh1^ before CDK2-mediated Cdh1 phosphorylation and inhibition. Future work mutating Wee1 and CDK phospho- sites in Cdh1 and identifying tyrosine phosphatases that act on Cdh1 will reveal the relative importance of each pathway for S-phase entry. Recent work showed that mTOR-mediated phosphorylation of Thr129 in Cdh1 transiently inactivated APC/C^Cdh1^, integrating metabolism with cell cycle entry^60^. The fact that this site is adjacent to Cdh1 Tyr130 phosphorylated by Wee1, and that phosphorylation of Ser131 has also been detected in cells ^59^, suggests a key role for this motif in regulating APC/C^Cdh1^ activity, most likely by stabilising the interaction of the C-motif in Cdh1 with Apc8. mTOR inhibition decreased Thr129 and Tyr130 phosphorylation in cells, yet mTOR was only shown to phosphorylate Thr129 ^60^. Therefore, mTOR phosphorylation may precede Wee1 phosphorylation of Cdh1 Tyr130. More broadly, our work, alongside others ^32,60,70,71^, suggests that APC/C^Cdh1^ is acting as a key signalling hub in G1, integrating signals from multiple inputs to determine whether cells commit to cell proliferation.

To date, CDK1 and CDK2 have been the only well characterised cell cycle substrates of Wee1 kinase. We now identify Cdh1 as a new Wee1 substrate. Wee1 can also phosphorylate Tyr37 in Histone H2B to reduce the transcription of histone genes ^72^. Recent work also suggested, but needs confirming, that TIMELESS/Tipin proteins involved in DNA replication may also be substrates of Wee1, highlighting that we have likely under-appreciated the substrates and functions of Wee1 ^73^.

Our experimental data is most consistent with the model that Wee1 inhibition of APC/C^Cdh1^ activity is independent of CDK2-mediated APC/C^Cdh1^ inhibition. While our experimental data are largely consistent with this model, there are two discrepancies. The first is that while both the model and experimental data show reduced Cdc25A during G1 after Wee1 inhibition, the model predicts that Cdc25A levels should increase after S-phase entry, due to APC/C^Cdh1^ being inactivated at the G1/S transition. This post-G1/S increase in Cdc25A does not occur in cells. This is most likely because Cdc25A is degraded by a second pathway in S-phase cells, driven by Chk1-mediated Cdc25A phosphorylation ^74,75^. Even in unperturbed cells, we see that Chk1 is required for Cdc25A degradation during S-phase since Chk1 inhibition leads to an increase in Cdc25A expression specifically in S-phase (Figure S10A). The second is that while our model is consistent with our data in that at very low levels of Wee1 activity, cells remain arrested in G0/G1, the model predicts high CDK2 activity and inhibition of Rb during the arrest, which is something we do not observe (Figure S10B). The reasons for this discrepancy are unclear but could be due to reduced CAK activity in cells that remain arrested ^76^.

Our data show that while Wee1i leads to increased CDK activity in cells, increased p21 expression generated downstream of Wee1i-induced DNA damage and premature inhibition of p21-degradation in S-phase, leads to G2 arrest. This is like what has been described in head and neck squamous carcinoma cells (HNSCC) treated with Wee1i ^77^ and explains previous observations demonstrating a role for p21 in protecting cells in S-phase from Wee1i-induced DNA damage ^78^. It is worth remembering that Wee1i were designed to be synthetically lethal with *TP53* mutations, where p53 induction of p21 expression downstream of DNA damage would be inactive. However, Wee1 inhibitors have efficacy in *TP53* wild-type tumours, demonstrating a need to understand the parameters that define Wee1i sensitivity ^79^. Here, we have identified the expression of p21 and APC/C^Cdh1^ complex components as potential indicators of sensitivity to Wee1i. We identify a potential use of Wee1i in treating *CDKN1A* mutant bladder cancer, which requires further investigation in pre- clinical models. Our data also suggest that Wee1i toxicity could emerge because of increased APC/C^Cdh1^ activity in quiescent stem cells that would impair their ability to enter proliferation and maintain tissue function.

## Methods

### Cell culture

All cell lines used are listed in Supplementary table 2.

#### Growth media

hTert-RPE1 and MCF7 cells were cultured at 37°C and 5% CO_2_ with the corresponding growth conditions: DMEM (Gibco, 11594486) supplemented with 10% Foetal Bovine Serum (FBS; Gibco F7514), 1% Penicillin-Streptomycin (P/S; Gibco 15140) while KU1919 and HBLAK were cultured in the same environmental conditions except that the media were RPMI 1640 (Gibco, A1049101) and Cnt Prime (Cellntec, CnT-PR), respectively.

#### Cell maintenance

All cell lines were routinely tested for mycoplasma using an Enzyme-Linked Immunosorbent Assay (ELISA) test kit (MycoAlert Cat no. LT07-318). Cells in culture were routinely split 1:10 every 3-4 days. For cell splitting, cells were washed with PBS and detached by incubating for 3 mins with Gibco Trypsin-EDTA (0.25%), phenol red (Gibco; 25200056) before being re-plated in fresh media. Accutase (ThermoFisher Scientific, 00-4555-56) was used for HBLAK.

### Cell synchronisation

#### Contact inhibition

RPE1 cells were arrested in G0/G1 by contact inhibition so that they reached 100% confluency for 3 days in a 6-well plate. After 6 days, cells were re-plated at low cell density in DMEM serum-free medium for 24 h. Cells were released into the cell cycle with addition of serum-containing DMEM (as above for growth medium).

#### CDK4/6i

RPE1 cells were treated with 0.5-1 µM Palbociclib (Selleckchem, S4482) for 24 h. Cells were washed three times with PBS to release cells back into the cell cycle with the addition of full growth medium. KU1919 cells were treated with 1 µM Palbociclib for 24 h. Cells were washed three times with PBS to release cells back into the cell cycle with the addition of full growth medium. MCF7 cells were treated with 1 µM Palbociclib for 24h. Cells were washed three times with PBS before release into the cell cycle with addition of appropriate media plus serum.

#### E. coli transformation

Chemically competent *E. coli* DH5a cells (NEB, C2987H) were used for transformation. Competent cells were thawed on ice for 30 minutes, after which 0.5μg of plasmid DNA of interest was added and incubated on ice for a further 20 minutes. To facilitate the uptake of plasmid DNA by the cells, they were subject to a heatshock for 45 seconds at 42°C in a water bath and subsequent incubation on ice for 2 minutes. After this, 250μL of room temperature S.O.C. medium was added to the cells and the mixture was incubated for an hour in a horizontal shaker at 37°C for 1 h. The bacteria were then spread on the appropriate antibiotic-containing LB agar plates and incubated at 37°C overnight.

### Cell line generation

#### Generation of endogenous Wee1-GFP-AID tagged hTert-RPE1 cell line

The GFP-AID-P2A-Neo tag (gift from Francis Barr’s lab^80^) was inserted at the C-terminus of the endogenous *WEE1* coding sequence using CRISPR/Cas9-mediated editing.

<u>gRNA cloning:</u> The gRNA used was forward 5’-ACCTCCCCTGAACACTGTG-3’, reverse 5’-CACAGTGTTCAGGGGGAGGT-3’. This gRNA sequence was cloned into the pX459 pSpCas9 using Bbsl restriction enzyme. pSpCas9(BB)-2A-Puro (PX459) V2.0 was a gift from Feng Zhang (Addgene plasmid 62988)^81^.

<u>Homology donor cloning:</u> The genomic region surrounding *WEE1* exon 11 (C-terminus) was PCR amplified from hTert-RPE1 gDNA using forward primer 5’- agtatttggcctattaaatttgtgttg-3’ and reverse primer 5’-tgcaaattgaagtgataatgttatttc-3’. *WEE1* exon 11 was cloned into pJet-vector (ThermoFisher) and the sequence verified by Sanger sequencing. To construct the homology donor plasmid, *WEE1* homology arms were PCR amplified from pJet-Wee1-Exon 11 using the high-fidelity Q5 DNA polymerase (NEB), according to the manufacturer’s instructions. To PCR amplify the Left Homology Arm (LHA), the following primers were used (the 5’ overhangs for Gibson assembly are underlined): forward primer: 5’- <u>acggtatcgataagcttgat</u>attcttttgatttattgcag-3’ and reverse primer: 5’- <u>gccccaccaccgctccacc</u>gtatatatagtaaggctgacag-3’ were used. To PCR amplify the Right Homology Arm (RHA), forward primer 5’-<u>cttcttgacgagttcttctgag</u>actactcctttc**a**cacctcc-3’ (in bold is the base mutated to change the PAM site in the recombined gene) and the reverse primer: 5’-<u>ccgggctgcaggaattcgat</u>gataaaaatatattttgtg-3’ were used. GFP-AID- GSG-P2A-Neo was PCR amplified from plasmid pFB8580 (generously provided by Francis Barr’s lab^80^) using the forward primer 5’-ggtgggagcggtggtggcg-3’ and reverse primer 5’-tcagaagaactcgtcaagaaggcgat-3’. The pBluescript (pSKII) vector was digested with EcoRV.

All PCR products were run on 0.8% TBE agarose gels, bands were gel extracted and assembled into EcoRV-digested pSKII(-) using Gibson Assembly with the NEB HiFi Assembly kit, according to the manufacturer’s instructions for four-way assembly. The final homology donor plasmid and the pX459 plasmid containing the gRNA were sequenced using Sanger sequencing to confirm the correct sequence.

<u>Generating Wee1-GFP-AID tagged cell line</u>: To generate the tagged cell lines, hTert- RPE1 myc-OsTIR1 cells (expressing a doxycycline-inducible myc-OsTir1 construct; a generous gift from Helfrid Hochegger^82^) were seeded in a 6 well plate at 5x10^4^ cells per well in 1.5 ml of growth media, one day before transfection. The next day, cells were forward transfected with Lipofectamine 2000 and 500ng of pX459-gRNA and 500ng of pSK-Wee1-GFP-AID-P2A-Neo. Cells were allowed to recover for 24 h before trypsinisation and seeding into 15 cm plate with fresh growth media. The next day, 1 mg/ml of G418 (ThermoFisher) was added to the plate, then fresh media plus G418 is added to the plate every 3-4 days until single colonies became visible. Single-cell colonies were picked using cloning cylinders (Sigma). Positives clones were first identified by nuclear GFP fluorescence before further analysis using western-blot.

#### Generation of endogenous Cdc25A-GFP tagged hTert-RPE1 cell line

gRNA cloning: To generate the gRNA targeting the C-terminus of human Cdc25A for CRISPR/Cas9-mediated gene tagging, gRNA forward primer: 5’- caccgaagaagctctgagggcggc-3’ and gRNA reverse primer: 5’- aaacgccgccctcagagcttcttc-3’ were annealed and ligated into BbsI-digested pSpCas9(BB)-2A-Puro (PX459) V2.0.

<u>Homology donor cloning:</u> To PCR amplify the *CDC25A* left and right homology arms from hTert-RPE1 genomic DNA, we used NEB Q5 High fidelity DNA polymerase and the following primers: for the left homology arm (the 5’ overhangs for Gibson assembly are underlined): forward 5’-<u>acggtatcgataagcttgat</u>gtgaaggtgctgggcaggactc-3’ and reverse 5’-<u>cgccaccaccgctcccacc</u>gagcttcttcagacgactgtac. For the right homology arm (in bold is the base mutated to remove the PAM sequence in recombined DNA): forward 5’-<u>atcgccttcttgacgagttcttctga</u>gggcggca**t**gaccagccagcag and reverse 5’- <u>ccgggctgcaggaattcgat</u>cagagcacctagctttctgtcc. GFP-GSG-P2A-Neo was PCR amplified from plasmid pFB8606 (generously provided by Francis Barr’s lab^80^), using the following primers: P2A-CT linker forward 5’-ggtgggagcggtggtggcg and P2A-CT Neo reverse 5’-tcagaagaactcgtcaagaaggcgat.

All PCR products were run on 0.8% TBE agarose gels, bands were gel extracted and assembled into EcoRV-digested pSKII(-) using Gibson Assembly with the NEB HiFi Assembly kit, according to the manufacturer’s instructions for four-way assembly. The final homology donor plasmid and the pX459 plasmid containing the gRNA were sequenced using Sanger sequencing to confirm the correct sequence.

<u>Generating Cdc25A-GFP tagged cell line</u>: hTert-RPE1 mRuby-PCNA cells in a 6-well plate (Sarstedt, 83.3920) were forward transfected with lipofectamine 2000 and 500ng of pX459-gRNA and 500ng of pSK-Cdc25A-CT-GFP, according to the manufacturer’s protocol, and left to recover overnight. The next day, cells were split into 15 cm tissue culture dishes (Corning, 420599) and 24h later, clones were selected with 1mg/ml G418. Media and antibiotic were refreshed every 3-4 days and after two weeks, single colonies were selected using cloning cylinders (Sigma) to transfer colonies to 24 well plates. Cells were expanded and initially screened for GFP positivity in the nucleus. gDNA was extracted from cells with GFP positive nuclei, using the Qiagen DNeasy blood and tissue kit, according to the manufacturer’s instructions. To check for targeted integration of the C-terminal tagging construct, the following primers were used: 5’-atgtacagtcgtctgaag and 5’-taccacagcgcctcctgg and all PCR product were sequenced by Sanger sequencing to check for in-frame tagging. Clones were then further validated by western blotting, growth curves and single-cell imaging to check cell cycle phase lengths.

#### PCNA tagging in hTert-RPE1 myc-OsTir1 Wee1-GFP-AID-Neo cells

To endogenously express the PCNA reporter, the fluorescent protein mRuby was inserted in the first exon into one allele of the PCNA locus by recombinant adeno-associated virus mediated (rAAV) homologous recombination^26^ in the hTert-RPE1 myc-OsTir1 Wee1-GFP-AID-Neo cell line. 293T cells were seeded in a T-75 Flask so that they reach 50-70% confluence at the time of transfection. 3 μg of plasmids pAAV-mRuby-PCNA (generous gift from Joerg Mansfeld, ICR), pRC and pHelper (9 μg total) were used and mixed with 3.75 ml of Optimem (Invitrogen). 19 μl of PLUS Reagent (Invitrogen) was added and mix was incubated 5-10 min at RT. 47 μl of Lipofectamine LTX (Invitrogen) was added to the tube and incubated for 30 min. 15 min before the end of the incubation, 293T cells were washed twice with PBS, then 7.5ml of Optimem (Invitrogen) was added and incubated at 37°C for 15 min. 1.5 ml of DNA/lipid mixture was added onto the cells and incubated at 37°C for 4-6 hours and then medium was replaced with complete DMEM. Two to three days post-infection, cells in medium (and any floating cells) were transferred to a sterile 50 ml Falcon tube. 5 ml of fresh medium was added to the flask and adherent cells were scraped off and transferred to the same 50 ml Falcon tube. Cells were lysed with three rounds of freeze-thaw (10min in dry ice/EtOH + 10 min in 37°C water bath). After each cycle, cells were vortexed at full speed for 1 min. Cells were then centrifuged at 10.000 x g for 30 min at 4°C. Tubes were wiped with EtOH and placed in the tissue culture hood. Cell-free supernatants were collected and rAAV preparations were dispensed in small aliquots (3-5 ml). hTert- RPE1 myc-OsTir1 Wee1-GFP-AID-Neo cells were seeded in a T-75 flask so that they reached 30-40% confluence at the time of infection. The T-75 flask was washed twice with PBS, then 5 ml of complete medium and 5 ml of rAAV stock were added and left to incubate at 37°C for 4 h. Fresh media was added to bring the volume up to 15 ml (or more) and infection was allowed to continue for 48h. After three days, mRuby-PCNA positive cells were sorted to single-cell density in 50:50 conditioned:growth media in 96-well plates by FACS. Cells were left to expand for two weeks and mRuby-PCNA positive clones were identified by microscopy.

#### Generation of p21 knockout cell lines

p21 knockout MCF7, KU1919 and HBLAK cell lines were generated as previously described^25^. The only modification was that HBLAK cells were transfected by nucleofection on the Amaxa nucleofector, using the MCF7 protocol. Knockouts were validated by immunostaining and western blotting for p21 after Nutlin-3 treatment, to boost the p21 signal.

### siRNA transfection

All siRNAs used are listed in Supplementary table 5.

For live imaging and western blotting experiments, cells were reverse transfected with siRNA in 384 well Phenoplates (Revvity) and 24-well plates (Greiner Bio-One, Cat. No. 662165), respectively. For imaging experiments, transfection was carried out 24h after CDK4/6 inhibition in a total volume of 30µL of media and used at a final concentration of 20 nM (10 nM for Cdh1 siRNA). The transfection was performed using Lipofectamine RNAiMax (Invitrogen 13778150), according to manufacturer’s instructions. In brief, 40 nL of Lipofectamine RNAiMax was added to 40 nL of 20 µM siRNA in 10 µL OptiMEM (Gibco 31985062) and 10 µL of the mixture was subsequently added to cells. Following transfection, cells were incubated 6h-24h, depending on the experiment. For western blotting, the same protocol was carried out with the following differences: cells were grown in 500 µL of media, 1µL of Lipofectamine RNAiMax was added to 1µL of 20 µM siRNA in 100µL OptiMEM and subsequently added to cells. Following transfection, cells were incubated 6h-24h, depending on the experiment.

### Immunostaining

All antibodies used for immunostaining are listed in Supplementary table 6.

Cells were grown in 96 or 384 well PhenoPlates (Revvity, 6055302 or 6057302, respectively). EdU was added to a final concentration of 10 μM for either 12h-24h at G0 release or with a 30 min pulse prior to fixation to identify to identify G0 cells and S-phase cells, respectively. Cells were fixed in 4% Formaldehyde/PBS for 15 min at RT, permeabilized with PBS/0.5% Triton X-100 for 15 min and blocked for 1 h in blocking buffer (PBS/2% BSA, filtered). Primary antibodies were diluted in blocking buffer and incubated overnight at 4 °C or for 2h at room temperature. After three washes in PBS, cells were incubated for 1h at RT in the dark in conjugated-secondary antibodies (1:1000; Invitrogen). For EdU incorporation, cells were incubated with 100 mM Tris-HCl pH 7.5, 4 mM CuS0_4_, 100 mM ascorbic acid and 5 μM sulfo-cyanine-5 azide for 30 min at RT in the dark, washed three times in PBS and counterstained with 1 μg/ml Hoechst for 10 min followed by three washes in PBS. All plate-washing steps were performed on an automated 50TS microplate washer (Biotek). Plates were imaged using an Operetta CLS (Revvity) with a 20X (N.A. 0.8) objective.

### Automated image analysis of fixed cells

All automated image analysis of Operetta CLS acquired images was performed using Harmony software (Revvity). Nuclei were segmented based on Hoechst intensity with nuclei at the edge of the field of view excluded from the analysis. Nuclear intensity of proteins was calculated as the mean intensity in the segmented region.

#### G0 calculation

G0 cells were identified by the absence of EdU staining. Briefly a nuclear to cytoplasmic (N:C) ratio was calculated by creating a four-pixel width ring around the segmented nucleus, and the ring intensity calculated as a proxy for the cytoplasmic portion. N:C ratios for individual cells were calculated and plotted in Prism (Graphpad). Cells with EdU N:C ratios <1.2 were called as G0.

### Western Blot

All antibodies used for western blotting are listed in Supplementary table 5.

Whole cell lysates were prepared following aspiration of media from culture plates, followed by washing with PBS on ice. Lysates were collected in 1X Novex Tris-Glycine-SDS sample buffer (Novex, LC2676) supplemented with 1x phosphatase inhibitors (ThermoScientific, 1862495), 1x protease inhibitors (ThermoScientific, 78429) and 1 mM DTT (BioUltra, 43816-10 ML). Samples were incubated at 95 °C for 10 min then centrifuged at 14,000 × *g* for 1 min before loading on Novex 4–20% Tris-Glycine gel (Invitrogen, XP04205). After transfer to PVDF-FL (Merck Life Sciences, IPFL00010), membranes were blocked in blocking buffer (TBS, 5% milk, 10% glycerol, Tween 0.1%) for 1 hr at RT and incubated in primary antibodies diluted in blocking buffer, overnight at 4 °C. Membranes were washed three times in TBS/0.05% TritonX- 100 before incubating for 1h at RT with HRP-conjugated secondary antibodies, diluted in blocking buffer. Membranes were washed three times in TBS/0.05% TritonX-100 and developed using Clarity Western ECL Substrate (Bio-Rad, 1705061) unless secondary antibodies antibodies were conjugated to Alexa Fluor. Blots were imaged on an Amersham Imager 680.

### Dox-inducible Wee1 degradation

For live cell imaging involving Wee1 degradation with Dox/IAA, RPE-1 Wee1-GFP-AID mRuby-PCNA cells were plated in 384-well Phenoplates (Revvity) at a density of 500 cells/well in 20 µL. Cells were synchronized with 1µM CDK4/6i for 24h before Dox/IAA (Doxycycline: 1µg/mL; IAA: 500µM) was added in the growth media when cells were released. For western blots, Dox/IAA was also added at release.

### *In vitro* kinase assay

All purified proteins used for in vitro kinase assays are listed in Supplementary table 7.

To confirm the activity of the Wee1 kinase we carried out a positive control experiment with CyclinB1/CDK1. Many thanks to Jon Pines (ICR) for the *in vitro* kinase protocol. For this reaction, 100 ng of recombinant CDK1-Cyclin B1 (Abcam) was diluted in 5 µL of kinase buffer (200 mM Tris pH 7.5, 150 mM NaCl, 10 mM MgCl2, 1 mM DTT) with 0.5 mM ATP and 18 µM RO-3306 (CDK1 inhibitor, Selleckchem). This mixture was added to 50 ng of recombinant Wee1 (ThermoFisher) diluted in 5 µL of kinase buffer with 0.5 µM MK-1775 (Wee1 inhibitor, Selleckchem) for the negative control. A negative control without Wee1 kinase addition was also carried out. The reactions were incubated for 30 min at 30°C and the reaction was stopped with addition of 5 µL of 3x SDS buffer with 100 mM DTT and boiled at 95°C for 5 min. Samples were then used for western blotting as described, with the exception that milk was replaced by BSA at the blocking step. pY15-CDK1/2/3/5 and CDK1 antibodies were used to quantify pY15-CDK1 phosphorylation. To monitor Cdh1 phosphorylation on tyrosine residues by Wee1, 80 ng of recombinant Strep-Cdh1 (see below) were diluted in 5 µL of kinase buffer with 0.5 mM ATP. This mixture was added to 50 ng of recombinant Wee1 diluted in 5 µL of kinase buffer with 0.5 µM MK-1775 for the negative control. A negative control without kinase addition was also carried out. The reactions were incubated for 30 min at 30°C and the reaction was stopped with addition of 5 µL of 3x SDS buffer with 100 mM DTT and boiled at 95°C for 5 min. Similar to the CDK1-Cyclin B1 reaction, samples were analyzed by western blotting. The pan-tyrosine antibody pTyr-100 and Cdh1 antibody were used to quantify the phosphorylation of tyrosine residues on Cdh1.

### Expression and purification of Strep-Cdh1

Expression and purification of Strep-Cdh1 was similar to that performed in^83^, with some modifications. Specifically, codon-optimised cDNAs of Strep-Cdh1 in pFastBac1 vectors were ordered from GeneArt (Thermo Fischer Scientific) and transformed into MultiBac cells for expression in insect cels. The baculovirus was created in Sf9 cells, this virus was then used to infect Hi5 cells at a cell density of 1 × 10^6^ cells/mL and then incubated for 72 hours at 27 °C, 130 rpm. All purification steps were carried out at 4 °C, first the cell pellet was thawed on ice in wash buffer (50 mM Tris-HCl pH 7.3, 500 mM NaCl, 5% glycerol, 0.5 mM TCE)] supplemented with 5 units/mL Benzonase and an EDTA-free protease inhibitor (Roche). The cells were sonicated and then centrifuged for 1 h at 48,000 × *g.* The supernatant was then bound to a 5 mL StrepTactin Superflow Plus cartridge (Qiagen) using a 1 mL/min flow rate. The column was washed extensively with the wash buffer and then eluted using elution buffer (50 mM Tris-HCl pH 7.0, 200 mM NaCl, 5 % glycerol, 0.5 mM TCEP, 2.5 mM Desthiobiotin).

### Proteomics

To determine the tyrosine residues phosphorylated by Wee1 on Cdh1, the reactions (5 replicates per condition) were carried out as described in the *in vitro* assay section, with the exception that the reactions were stopped with addition of 1M Guanidine hydrochloride. We diluted 5 μL of the kinase reaction with 45 μL water containing TCEP (final concentration 10 mM) and chloracetamide (20 mM). The reaction was incubated at 37 °C to reduce and alkylate Cys and digestion was initiated by addition of trypsin and LysC at 5 and 10 ng/µL final concentration respectively. After overnight digestion the samples were desalted on 3mg of oasis HLB buffer. The samples were evaporated in a speed vac, resuspended in 30 µL of 0.1% TFA and 5 µL analysed by LC-MS.

Chromatographic separation was performed using an Ultimate 3000 RSLC nano liquid chromatography system coupled to a Exploris 240 mass spectrometer via an EASY-Spray source. Electro-spray ionisation was achieved with Bruker PepSep emitters (PN: PSFSELJ10, 10µm). Peptide solutions were injected directly onto the analytical column (self-packed column, CSH C18 1.7µm beads, 150μm × 30cm) at a working flow rate of 1.25 μL/min for 4 minutes. Peptides were then separated using a 60-minute stepped gradient: 0-45% of buffer B for 70 minutes (composition of buffer A – 95/5%: H2O/DMSO + 0.1% FA, buffer B – 75/20/5% MeCN/H2O/DMSO + 0.1% FA), followed by column conditioning and equilibration. Eluted peptides were analysed in DIA mode: an initial MS1 scan was carried out at 140,000 resolution with an AGC target of 3e6 for a maximum IT of 200ms, m/z range: 410-1600. This was followed by thirty 30K resolution MS2 scans covering 410-1600 with variable windows sizes predicted by encyclopeDIA. AGC target was set to 3e6 with maximum IT on auto. Normalised collision energy was set to 27%.

Data were processed using the Spectronaut software platform (Biognosys, 19.9.250512.62635) in direct DIA mode. Searches were carried out against *S.frugiperda* concatenated with human Wee1 sequence, *T.Ni* protein sequence database concatenated with human Cdh1 sequence and a universal protein contaminants database ^84^. The DIA analysis was performed with higher stringency settings than in the default workflow as described previously ^85^. Default settings were used for PTM analysis with probability cut-off for PTM localization set to 0, prior to table export. To account for Cdh1 intensity variance at protein level, phospho-site intensities were also normalised to the total protein level. Specific sites of interest were validated using Spectronaut feature viewer.

The two idenitifed phosphosites were located on DGLAY[Phospho (Y)]SALLK – Y91 GLFTY[Phospho (Y)]SLSTK - Y130 peptides both with 2+ charge. We then reran the samples targeting the peptides in PRM and SIM modes at 240K resolution and 500 ms maximum injection time.

The mass spectrometry proteomics data have been deposited to the ProteomeXchange Consortium via the PRIDE ^86^ partner repository with the dataset identifier PXD068577.

### Fluorescence-activated cell sorting

Fluorescence-activated cell sorting of cells was performed at the Flow cytometry facility, MRC Laboratory of Medical Sciences (MRC LMS). Cells were washed with PBS and resuspended in FACS sorting buffer (1 ml of PBS 2% FBS) to a concentration of 8-12 million cells/ml. Prior to sorting, cells were filtered through a 30-40 μM filter.

### Growth curves

Cells were plated at a density of 1000 cells per well in triplicate in 96 well tissue culture plates. Brightfield images were taken every 4 h for 7 days using the 10x (N.A. 0.3) objective and percent confluency calculated on the CellCyteX (Echo) live imaging system. To calculate confluency, a mask was created using the default settings on the CellCyteX (brightness 0%, contrast 0%, contrast sensitivity 50 a.u., smoothing 2 a.u. and filled hole size 100 µm^2^). Results were plotted with python (Seaborn package).

### Drug dose curves – IC50

Cells were plated at a density of 500 cells/well in 20 µL. For the drug dose curve assays with siRNA transfection, cells were transfected (10 µL) for 6h prior to drug addition as described previously. MK-1775 was prepared as a 7x1:3 serial dilutions (7x1:4 in case of transfection) in growth media with final concentrations for all assays that were 100 µM, 10 µM, 1 µM, 0.1 µM, 0.01 µM, 0.001 µM. 10 µL were added per well. After 72h of incubation, cells were fixed and stained for Hoechst. Plates were imaged using an Operetta CLS (Revvity) with a 20X (N.A. 0.8) objective. Cell viability was calculated as a percentage and normalized IC_50_ values were computed using GraphPad Prism.

### Emi1 overexpression

hTert-RPE1 CCNB1-eYFP mTurq2-EMI1 53BP1-mCherry cells were (a generous gift from Rene Medema^87^) seeded in a 384-well Phenoplates (Revvity) at a density of 500 cells per well in 20 µL. Cells were arrested for 24h in CDK4/6i and then released with Doxycycline (1µg/mL) to induce Emi1 expression, EdU and DMSO or Wee1i for 12h prior to fixation to identify G0 cells. Cells were fixed in 4% Formaldehyde/PBS for 15 min at RT, permeabilized with PBS/0.5% Triton X-100 for 15 min and blocked for 1 h in blocking buffer (PBS/2% BSA, filtered). For EdU incorporation, cells were incubated with 100 mM Tris-HCl pH 7.5, 4 mM CuS0_4_, 100 mM ascorbic acid and 5 μM sulfo-cyanine-5 azide for 30 min at RT in the dark, washed three times in PBS and counterstained with 1 μg/ml Hoechst for 10 min followed by three washes in PBS. The inhibition of Cdh1 was confirmed by western blot. Cells were plated at a density of 40000 cells per well in a 24-well plate, then arrested for 24 hr with CDK4/6i and released with addition of doxycycline to induce Emi1 expression (1 and 10 µg/mL) for 4 hours before cell lysis in 1X Novex Tris-Glycine-SDS sample buffer (Novex, LC2676) supplemented with 1x phosphatase inhibitors (ThermoScientific, 1862495), 1x protease inhibitors (ThermoScientific, 78429) and 1 mM DTT (BioUltra, 43816-10 ML).

### Timelapse live imaging

Live cell imaging was performed on the Operetta CLS High-Content Analysis microscope (Revvity), with atmospheric control to maintain cells at 37°C, 5% CO2 and with a breathable membrane (Thermofisher) to stop evaporation. Cells were plated in a 384-well Phenoplates (Revvity) at a density of 500 cells/well in 20μL of media for 24 h (unless otherwise stated) prior to imaging. Cells were imaged using a 20X (N.A. 0.8) objective at 10-min intervals for 48 h (unless otherwise stated) in 50μL phenol red-free media. For hTert-RPE1 Wee1-GFP-AID mRuby-PCNA and hTert-RPE1 mRuby-PCNA Cdc25A-GFP live imaging, cells were imaged on an automated inverted spinning disk confocal microscope system IX83 (Olympus) combined with CellVivo (Olympus). Cells were plated on IbiTreat μ-slide 8well (ibidi) plates at a density of 5000 cells/well in 200μL of media or in a 384-well Phenoplates (Revvity) plates at a density of 500 cells/well in 20μL of media, 24 h prior to imaging (unless otherwise stated). Images were typically acquired with a 20x plan (UCPLFLN) fluorescence objective (NA 0.7).

### Automated image analysis of live imaging experiments

#### Automated cell tracking (Nuclitrack)

Images were exported as tiff files and analysis was performed using NucliTrack^88^ and/or FIJI and Napari ^89^. All cells analysed were present throughout the entire movie unless they died. Spontaneous death was quantified manually. mRuby-PCNA was used to define cell cycle phases as described previously^25,26^.

#### Automated cell tracking (Cellpose + Ultrack)

The code used for single-cell time-lapse image segmentation and tracking was implemented in Python (jupyter notebook) and is available on a GitHub repository (https://github.com/Alexis-Barr-Lab). This analysis pipeline combines Cellpose^90^ segmentation and tracking using Ultrack ^91^. This analysis pipeline was run with the Jex HPC cluster at the MRC Laboratory of Medical Sciences ^92^. In Brief, raw TIFF stacks (shape: T × Y × X) were background-subtracted using a rolling-ball–style method via a Gaussian filter (σ = 15 px) followed by morphological reconstruction to separate foreground from low-frequency background. Frames were normalized (γ = 0.5) and stored in Zarr format for efficient chunked I/O. Normalized frames were processed with a Cellpose model (cpsam). Masks for each time point were generated, yielding per-frame 2D label arrays written into a Zarr volume. To recover fine cell boundaries and split merged objects, a watershed was run on each frame. Thresholding and binary morphology mask detections, followed by distance-transform watershed with minima suppression, generated second label sets, also saved as Zarr. GPU-accelerated boundary detection was implemented in CuPy; boundaries from both Cellpose and watershed masks were combined and smoothed (σ = 5) to create high-resolution contour maps for downstream linking. Centroids of watershed segments between frames were calculated. The Hungarian algorithm (linear_sum_assignment) linked cells, providing centroid-distance statistics: 95th percentile, maximum, average, and 25th percentile displacements – informing Ultrack’s max_distance parameter. Ultrack was then configured and executed. For quality checks, tracks and labels were exported and visualized. Background subtraction is then applied to the channel of interest and properties are exported.

## Statistical analysis

Statistical analysis used in this manuscript were performed in GraphPad Prism unless stated otherwise.

## Generation of MCF7 Wee1i resistant cells

In order to generate MCF7 Wee1i-resistant cells, MCF7 wild-type cells were plated onto a 15 cm tissue culture dish (Corning, 420599) at a density of 5 x 10^6^ cells/mL. Cells were treated with 0.075µM MK-1775 24 hours later and incubated until confluence. Cells were then split and treated with double amount of MK-1775 24 hr later and incubated until confluence. This step was repeated until 0.6 µM MK-1775. The resistance of these cells was measured against the parental cells with a 72 hrs drug dose curve (IC50).

## RNAseq

MCF7-Resistant and wild-type cells were plated at a density of 100000 cells per well in a 6-well plate (Sarstedt, 83.3920). 24 hr later, total RNA was extracted using an RNeasy MinElute Cleanup Kit (Qiagen 74204) according to manufacturer’s instructions. Quality and concentration were assessed using the Agilent 2100 Bioanalyser RNA 6000 Nano assay.

RNAseq libraries were made from 200ng of Total RNA using the Watchmaker Genomics mRNA enrichment and RNA Library Prep Kits according to manufacturer’s instructions. Libraries were dual indexed using IDT xGen Unique Dual Index Primers and library quality was assessed using the Agilent 2100 Bioanalyser High-Sensitivity DNA assay. Libraries were pooled and run on an Illumina NextSeq2000, generating a minimum of 60 million Paired End 60bp reads per sample.

RNA-seq reads were processed using cutadapt v4.7^93^ to remove Illumina adapters, to quality trim at Q20, and to filter read pairs containing N bases or where either read <31 bps. Processed reads were quantified with Salmon v1.9.0^94^ using transcripts from the GRCh38 Ensembl v107 annotations. Salmon’s expectation maximisation procedures were set to enable modelling of sequencing and GC biases.

Within R, transcript level counts from Salmon were imported and aggregated to the gene level using tximport v1.32.0^95^. Sample quality assessments were performed in relation to collected RNA integrity (RIN) values, sample concentrations used in library preparation (ng/ul), read GC content, and percentage read mapping. Normalisation, further PCA- and clustering-based quality control based on normalised values, and differential expression analyses were performed with DESeq2 v1.44.0^96^. Independent hypothesis weighting was conducted to optimise the power of p-value filtering^97^ using IHW v1.32.0 and log2 fold change shrinkage was performed using ashr v2.2-63^98^ to reduce the impact of low expression on estimation of fold change values. Multiple correction testing adjustments were optimised by passing thresholds for both false discovery rate and log2 fold change at the point of results generation within DESeq2. Significance was assessed using a *q* value of 0.01.

## Cycloheximide Cdh1 protein stability assay

To monitor Cdh1 stability upon Wee1 inhibition, RPE1 mRuby-PCNA cells were plated in a 24-well plate at a density of 50000 cells per well. Cells were arrested with CDK4/6i for 24 hrs and treated with Wee1i (1µM) or DMSO and Cycloheximide (100 µg/mL) upon release. Subsequently, cells were lysed every hour for 4 hrs in 1X Novex Tris-Glycine-SDS sample buffer (Novex, LC2676) supplemented with 1x phosphatase inhibitors (ThermoScientific, 1862495), 1x protease inhibitors (ThermoScientific, 78429) and 1 mM DTT (BioUltra, 43816-10 ML).

## APC/C activity calculation

APC/C activity (*k*_APC_) calculations based on mCherry-Geminin and Cyclin A2-mVenus levels (APC degron) were calculated as previously described^32,99^ with equation (1).

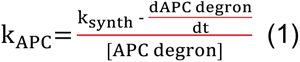

## Protein alignment

Clustal Omega^100^ was used to align Cdh1 protein sequences and outputs (Consensus frequency of residues, Identity matrix and phylogenetic tree) were plotted in python.

## Modelling

We built our cell cycle model using a system of coupled nonlinear differential equations describing the rates of change of biochemical species responsible for the regulation of the G1 and S states. The regulation of subsequent cell cycle phases was omitted for simplicity, as our experimental approach focused on quiescence release, rather than cycling cells. The description of the modelling and as well as the parameters used are available in the Supplementary material.

## Data accessibility

All RNAseq data generated in this study are publicly available in Gene Expression Omnibus (GEO) at GSE302212.

The mass spectrometry proteomics data have been deposited to the ProteomeXchange Consortium via the PRIDE partner repository with the dataset identifier PXD068577.

## Acknowledgements

We thank colleagues in the Barr lab for helpful discussions. We thank Rose Young and Claudio Alfieri (ICR) for providing purified Cdh1 protein. We thank Rene Medema (NKI) and Jean Cook (UNC Chapel Hill) for providing cell lines. We thank Jon Pines (ICR) for the Wee1 *in vitro* kinase protocol and Francis Barr (University of Oxford) for the homology donor plasmids for CRISPR/Cas9 gene-tagging. We acknowledge use of the Jex HPC cluster at MRC LMS and the resources and support provided by the IT and Bioinformatics facilities. We thank MRC LMS/NIHR Imperial Biomedical Research Centre Flow Cytometry Facility for support. We also thank Chad Whilding and MRC LMS core microscopy for their help and support. ARB, LAV and BRP were supported by a CRUK Career Development Fellowship to ARB (C63833/A25729), and work in the Barr lab was supported by MRC LMS core funding (MC-A658-5TY60).

## Author contributions

ARB supervised experiments, helped to analyse data, secured funding for the work and wrote the manuscript. LAV, LGM, BRP, EH, LCL, WAW, JEB and EG performed experiments and analysed data. CMD and BN developed mathematical models. LAV, LGM, CMD and BRP prepared figures and helped to write the manuscript. RY and CA provided Strep-Cdh1 for in vitro kinase assays. IA and LG prepared samples for RNA-seq analysis. GR and PS prepared samples for proteomics. AM and PS analysed mass spectrometry data. GY analysed RNA-seq data.

## Competing interests

The authors declare no conflict of interest.

## Materials and Correspondence

Please address all requests for materials to Alexis Barr

## Notes

### Competing Interest Statement

The authors have declared no competing interest.

## References

1. Parker, L. L. & Piwnica-Worms, H. Inactivation of the p34cdc2-Cyclin B Complex by the Human WEE1 Tyrosine Kinase . Science (1979) 257, 1955–1957 (1992).

2. Heald, R., McLoughlin, M. & McKeon, F. Human wee1 maintains mitotic timing by protecting the nucleus from cytoplasmically activated cdc2 kinase. Cell 74, 463–474 (1993).

3. McGowan, C. H. & Russell, P. Human Wee1 kinase inhibits cell division by phosphorylating p34cdc2 exclusively on Tyr15. EMBO J 12, 75–85 (1993).

4. Parker, L. L. & Piwnica-Worms, H. Inactivation of the p34cdc2-Cyclin B Complex by the Human WEE1 Tyrosine Kinase . Science (1979) 257, 1955–1957 (1992).

5. Gautier, J., Solomon, M. J., Booher, R. N., Bazan, J. F. & Kirschner, M. W. cdc25 is a specific tyrosine phosphatase that directly activates p34cdc2. Cell 67, 197– 211 (1991).

6. Kumagai, A. & Dunphy, W. G. The cdc25 protein controls tyrosine dephosphorylation of the cdc2 protein in a cell-free system. Cell 64, 903–914 (1991).

7. Strausfeld, U. et al. Dephosphorylation and activation of a p34cdc2/cyclin B complex in vitro by human CDC25 protein. Nature 351, 242–245 (1991).

8. Millar, J. B. A. et al. p55CDC25 is a nuclear protein required for the initiation of mitosis in human cells. Proc Natl Acad Sci U S A 88, 10500–10504 (1991).

9. Lammer, C. et al. The cdc25B phosphatase is essential for the G 2/M phase transition in human cells. J Cell Sci 111, 2445–2453 (1998).

10. El-Deiry, W. S. et al. WAF1/CIP1 Is Induced in p53-mediated G1 Arrest and Apoptosis1. Cancer Res 54, 1169–1174 (1994).

11. Dulić, V. et al. p53-dependent inhibition of cyclin-dependent kinase activities in human fibroblasts during radiation-induced G1 arrest. Cell 76, (1994).

12. Di Leonardo, A., Linke, S. P., Clarkin, K. & Wahl, G. M. DNA damage triggers a prolonged p53-dependent G1 arrest and long-term induction of Cip1 in normal human fibroblasts. Genes Dev 8, (1994).

13. Hirai, H. et al. Small-molecule inhibition of Wee1 kinase by MK-1775 selectively sensitizes p53-deficient tumor cells to DNA-damaging agents. Mol Cancer Ther 8, 2992–3000 (2009).

14. Hirai, H. et al. MK-1775, a small molecule Wee1 inhibitor, enhances anti-tumor efficacy of various DNA-damaging agents, including 5-fluorouracil. Cancer Biol Ther 9, 514–522 (2010).

15. Bridges, K. A. et al. MK-1775, a Novel Wee1 Kinase Inhibitor, Radiosensitizes p53-Defective Human Tumor Cells. Clinical Cancer Research 17, 5638–5648 (2011).

16. Otto, T. & Sicinski, P. Cell cycle proteins as promising targets in cancer therapy. Nat Rev Cancer 17, 93–115 (2017).

17. Zhang, C., Peng, K., Liu, Q., Huang, Q. & Liu, T. Adavosertib and beyond: Biomarkers, drug combination and toxicity of WEE1 inhibitors. Crit Rev Oncol Hematol 193, 104233 (2024).

18. Hughes, B. T., Sidorova, J., Swanger, J., Monnat, R. J. & Clurman, B. E. Essential role for Cdk2 inhibitory phosphorylation during replication stress revealed by a human Cdk2 knockin mutation. Proceedings of the National Academy of Sciences 110, 8954–8959 (2013).

19. Zhao, H., Chen, X., Gurian-west, M. & Roberts, J. M. Phosphorylation in a CDK2AF Knock-In Mouse Causes Misregulation of DNA Replication and Centrosome Duplication. 2, 1421–1432 (2012).

20. Baldin, V., Cans, C., Watanabe, N. & Ducommun, B. Evidence for a Mammalian Nim1-like Kinase Pathway Acting at the G0-1/S Transition. Biochem Biophys Res Commun 236, 130–134 (1997).

21. Wu, C. L. et al. Cables enhances cdk2 tyrosine 15 phosphorylation by Wee1, inhibits cell growth, and is lost in many human colon and squamous cancers. Cancer Res 61, 7325–7332 (2001).

22. 22. Watanabe’, N., Broome, M. & Hunter, T. Regulation of the Human WEE1Hu CDK Tyrosine 15-Kinase during the Cell Cycle. The EMBO Journal vol. 14 https://www.ncbi.nlm.nih.gov/pmc/articles/PMC398287/pdf/emboj00033- 0044.pdf (1995).

23. Wu, C. L. et al. Cables enhances cdk2 tyrosine 15 phosphorylation by Wee1, inhibits cell growth, and is lost in many human colon and squamous cancers. Cancer Res 61, 7325–7332 (2001).

24. Barr, A. R., Heldt, F. S., Zhang, T., Bakal, C. & Novák, B. A Dynamical Framework for the All-or-None G1/S Transition. Cell Syst 2, (2016).

25. Barr, A. R. et al. DNA damage during S-phase mediates the proliferation- quiescence decision in the subsequent G1 via p21 expression. Nat Commun 8, (2017).

26. Zerjatke, T. et al. Quantitative Cell Cycle Analysis Based on an Endogenous All- in-One Reporter for Cell Tracking and Classification. Cell Rep 19, 1953–1966 (2017).

27. Trotter, E. W. & Hagan, I. M. Release from cell cycle arrest with Cdk4/6 inhibitors generates highly synchronized cell cycle progression in human cell culture: Cdk4/6 Induction Synchronisation. Open Biol 10, (2020).

28. Matson, J. P. et al. Intrinsic checkpoint deficiency during cell cycle re-entry from quiescence. Journal of Cell Biology 218, 2169–2184 (2019).

29. Spencer, S. L. et al. The Proliferation-Quiescence Decision Is Controlled by a Bifurcation in CDK2 Activity at Mitotic Exit. Cell 155, 369–383 (2013).

30. Pennycook, B. R. & Barr, A. R. Restriction point regulation at the crossroads between quiescence and cell proliferation. FEBS Lett (2020).

31. Sørensen, C. S. et al. A Conserved Cyclin-Binding Domain Determines Functional Interplay between Anaphase-Promoting Complex–Cdh1 and Cyclin A-Cdk2 during Cell Cycle Progression. Mol Cell Biol 21, 3692 (2001).

32. Cappell, S. D., Chung, M., Jaimovich, A., Spencer, S. L. & Meyer, T. Irreversible APCCdh1 Inactivation Underlies the Point of No Return for Cell-Cycle Entry. Cell 166, 167–180 (2016).

33. Lukas, C. et al. Accumulation of cyclin B1 requires E2F and cyclin-A-dependent rearrangement of the anaphase-promoting complex. Nature 1999 401:6755 401, 815–818 (1999).

34. Grant, G. D., Kedziora, K. M., Limas, J. C., Cook, J. G. & Purvis, J. E. Accurate delineation of cell cycle phase transitions in living cells with PIP-FUCCI. Cell Cycle 17, 2496–2516 (2018).

35. Geley, S. et al. Anaphase-Promoting Complex/Cyclosome–Dependent Proteolysis of Human Cyclin a Starts at the Beginning of Mitosis and Is Not Subject to the Spindle Assembly Checkpoint. J Cell Biol 153, 137 (2001).

36. Hua, X. H., Yan, H. & Newport, J. A role for Cdk2 kinase in negatively regulating DNA replication during S phase of the cell cycle. Journal of Cell Biology 137, 183–192 (1997).

37. Blow, J. J. & Hodgson, B. Replication licensing — Origin licensing: defining the proliferative state? Trends Cell Biol 12, 72–78 (2002).

38. Yan, Z. et al. Cdc6 is regulated by E2F and is essential for DNA replication in mammalian cells. Proc Natl Acad Sci U S A 95, 3603–3608 (1998).

39. Beck, H. et al. Cyclin-Dependent Kinase Suppression by WEE1 Kinase Protects the Genome through Control of Replication Initiation and Nucleotide Consumption. Mol Cell Biol 32, 4226 (2012).

40. Beck, H. et al. Regulators of cyclin-dependent kinases are crucial for maintaining genome integrity in S phase. Journal of Cell Biology 188, 629–638 (2010).

41. Elbæk, C. R. et al. WEE1 kinase protects the stability of stalled DNA replication forks by limiting CDK2 activity. Cell Rep 38, (2022).

42. Moldovan, G. L., Pfander, B. & Jentsch, S. PCNA, the Maestro of the Replication Fork. Cell 129, 665–679 (2007).

43. Nishitani, H. et al. CDK inhibitor p21 is degraded by a proliferating cell nuclear antigen-coupled Cul4-DDB1Cdt2 pathway during s phase and after UV irradiation. Journal of Biological Chemistry 283, (2008).

44. Kim, Y., Starostina, N. G. & Kipreos, E. T. The CRL4Cdt2 ubiquitin ligase targets the degradation of p21Cip1 to control replication licensing. Genes Dev 22, (2008).

45. Abbas, T. et al. PCNA-dependent regulation of p21 ubiquitylation and degradation via the CRL4Cdt2 ubiquitin ligase complex. Genes Dev 22, (2008).

46. Galanos, P. et al. Chronic p53-independent p21 expression causes genomic instability by deregulating replication licensing. Nat Cell Biol 10.1038/ncb3378 (2016) doi: 10.1038/ncb3378.

47. El-Deiry, W. S. et al. WAF1, a potential mediator of p53 tumor suppression. Cell 75, 817–825 (1993).

48. Moiseeva, T. N., Qian, C., Sugitani, N., Osmanbeyoglu, H. U. & Bakkenist, C. J. WEE1 kinase inhibitor AZD1775 induces CDK1 kinase-dependent origin firing in unperturbed G1- and S-phase cells. Proc Natl Acad Sci U S A 116, 23891– 23893 (2019).

49. Domínguez-Kelly, R. et al. Wee1 controls genomic stability during replication by regulating the Mus81-Eme1 endonuclease. Journal of Cell Biology 194, 567– 579 (2011).

50. Rizzardi, L. F. et al. CDK1-dependent Inhibition of the E3 Ubiquitin Ligase CRL4CDT2 Ensures Robust Transition from S Phase to Mitosis. Journal of Biological Chemistry 290, 556–567 (2015).

51. Vassilopoulos, A. et al. WEE1 murine deficiency induces hyper-activation of APC/C and results in genomic instability and carcinogenesis. Oncogene 34, 3023–3035 (2015).

52. Brandeis, M. & Hunt1, T. The proteolysis of mitotic cyclins in mammalian cells persists from the end of mitosis until the onset of S phase. EMBO J 15, 5280– 5289 (1996).

53. Mailand, N. & Diffley, J. F. X. CDKs Promote DNA Replication Origin Licensing in Human Cells by Protecting Cdc6 from APC/C-Dependent Proteolysis. Cell 122, 915–926 (2005).

54. Donzelli, M. et al. Dual mode of degradation of Cdc25 A phosphatase. EMBO J 21, 4875–4884 (2002).

55. Hoffmann, I., Draetta, G. & Karsenti, E. Activation of the phosphatase activity of human cdc25A by a cdk2-cyclin E dependent phosphorylation at the G1/S transition. EMBO J 13, 4302–4310 (1994).

56. Sigi, R. et al. Loss of the mammalian APC/C activator FZR1 shortens G1 and lengthens S phase but has little effect on exit from mitosis. J Cell Sci 122, 4208– 4217 (2009).

57. Hsu, J. Y., Reimann, J. D. R., Sørensen, C. S., Lukas, J. & Jackson, P. K. E2F- dependent accumulation of hEmi1 regulates S phase entry by inhibiting APCCdh1. Nat Cell Biol 4, (2002).

58. Feringa, F. M. et al. Hypersensitivity to DNA damage in antephase as a safeguard for genome stability. Nature Communications 2016 7:*1* **7**, 1–10 (2016).

59. Hornbeck, P. V. et al. PhosphoSitePlus, 2014: Mutations, PTMs and recalibrations. Nucleic Acids Res 43, (2015).

60. Paul, D. et al. Transient APC/C inactivation by mTOR boosts glycolysis during cell cycle entry. Nature 2025 1–10 (2025) doi:10.1038/s41586-025-09328-w.

61. Höfler, A. et al. Cryo-EM structures of apo-APC/C and APC/CCDH1:EMI1 complexes provide insights into APC/C regulation. Nat Commun 15, 10074 (2024).

62. Schwab, M., Neutzner, M., Möcker, D. & Seufert, W. Yeast Hct1 recognizes the mitotic cyclin Clb2 and other substrates of the ubiquitin ligase APC. EMBO J 20, 5165 (2001).

63. Vodermaier, H. C., Gieffers, C., Maurer-Stroh, S., Eisenhaber, F. & Peters, J. M. TPR subunits of the anaphase-promoting complex mediate binding to the activator protein CDH1. Current Biology 13, 1459–1468 (2003).

64. Zhang, J. et al. Predicting protein-protein interactions in the human proteome. Science (1979) 0, eadt1630 (2025).

65. Abbas, T. & Dutta, A. p21 in cancer: intricate networks and multiple activities. Nat Rev Cancer 9, 400–414 (2009).

66. Lawson, A. R. J. et al. Extensive heterogeneity in somatic mutation and selection in the human bladder. Science (1979) 370, 75–82 (2020).

67. O’Leary, B., Finn, R. S. & Turner, N. C. Treating cancer with selective CDK4/6 inhibitors. Nat Rev Clin Oncol 13, 417–430 (2016).

68. Hughes, B. T., Sidorova, J., Swanger, J., Monnat, R. J. & Clurman, B. E. Essential role for Cdk2 inhibitory phosphorylation during replication stress revealed by a human Cdk2 knockin mutation. Proceedings of the National Academy of Sciences 110, 8954–8959 (2013).

69. Zhao, H., Chen, X., Gurian-west, M. & Roberts, J. M. Phosphorylation in a CDK2AF Knock-In Mouse Causes Misregulation of DNA Replication and Centrosome Duplication. 2, 1421–1432 (2012).

70. Wan, L. et al. The APC/C E3 ligase complex activator fzr1 restricts braf oncogenic function. Cancer Discov 7, (2017).

71. Kernan, J., Bonacci, T. & Emanuele, M. J. Who guards the guardian? Mechanisms that restrain APC/C during the cell cycle. Biochimica et Biophysica Acta - Molecular Cell Research vol. 1865 Preprint at 10.1016/j.bbamcr.2018.09.011 (2018).

72. Mahajan, K., Fang, B., Koomen, J. M. & Mahajan, N. P. H2B Tyr37 phosphorylation suppresses expression of replication-dependent core histone genes. Nature Structural & Molecular Biology 2012 19:*9* **19**, 930–937 (2012).

73. Sebastian, R. et al. Mechanism for local attenuation of DNA replication at double-strand breaks. Nature 2025 639:8056 **639**, 1084–1092 (2025).

74. Mailand, N. et al. Rapid destruction of human Cdc25A in response to DNA damage. Science (1979) 288, (2000).

75. Goto, H. et al. Chk1-mediated Cdc25A degradation as a critical mechanism for normal cell cycle progression. J Cell Sci 10.1242/jcs.223123 (2019) doi: 10.1242/jcs.223123.

76. Hirschi, A. et al. Article A Cdk7-Cdk4 T-Loop Phosphorylation Cascade Promotes G1 Progression. 10.1016/j.molcel.2013.04.003 (2013) doi: 10.1016/j.molcel.2013.04.003.

77. Diab, A. et al. FOXM1 drives HPV+ HNSCC sensitivity to WEE1 inhibition. Proc Natl Acad Sci U S A 117, (2020).

78. Hauge, S., Macurek, L. & Syljuåsen, R. G. p21 limits S phase DNA damage caused by the Wee1 inhibitor MK1775. Cell Cycle 18, 834–847 (2019).

79. Van Linden, A. A. et al. Inhibition of Wee1 sensitizes cancer cells to anti- metabolite chemotherapeutics in vitro and in vivo, independent of p53 functionality. Mol Cancer Ther 12, 2675 (2013).

80. Alfonso-Pérez, T., Hayward, D., Holder, J., Gruneberg, U. & Barr, F. A. MAD1- dependent recruitment of CDK1-CCNB1 to kinetochores promotes spindle checkpoint signaling. Journal of Cell Biology 218, 1108–1117 (2019).

81. Ran, F. A. et al. Genome engineering using the CRISPR-Cas9 system. Nat Protoc 8, 2281 (2013).

82. Hégarat, N. et al. Cyclin A triggers Mitosis either via the Greatwall kinase pathway or Cyclin B. EMBO J 39, e104419 (2020).

83. Chang, L., Zhang, Z., Yang, J., McLaughlin, S. H. & Barford, D. Atomic structure of the APC/C and its mechanism of protein ubiquitination. Nature 2015 522:7557 **522**, 450–454 (2015).

84. Frankenfield, A. M., Ni, J., Ahmed, M. & Hao, L. Protein Contaminants Matter: Building Universal Protein Contaminant Libraries for DDA and DIA Proteomics. J Proteome Res 21, 2104–2113 (2022).

85. Baker, C. P., Bruderer, R., Abbott, J., Arthur, J. S. C. & Brenes, A. J. Optimizing Spectronaut Search Parameters to Improve Data Quality with Minimal Proteome Coverage Reductions in DIA Analyses of Heterogeneous Samples. J Proteome Res 23, 1926–1936 (2024).

86. Perez-Riverol, Y. et al. The PRIDE database at 20 years: 2025 update. Nucleic Acids Res 53, D543–D553 (2025).

87. Hornsveld, M. et al. A FOXO-dependent replication checkpoint restricts proliferation of damaged cells. Cell Rep 34, 108675 (2021).

88. Cooper, S., Barr, A. R., Glen, R. & Bakal, C. NucliTrack: An integrated nuclei tracking application. Bioinformatics 10.1093/bioinformatics/btx404(2017) doi: 10.1093/bioinformatics/btx404.

89. Sofroniew, N. et al. napari: a multi-dimensional image viewer for Python. 10.5281/ZENODO.17367124 doi: 10.5281/ZENODO.17367124.

90. Pachitariu, M., Rariden, M. & Stringer, C. Cellpose-SAM: superhuman generalization for cellular segmentation. bioRxiv 2025.04.28.651001 (2025) doi:10.1101/2025.04.28.651001.

91. Bragantini, J. et al. Ultrack: pushing the limits of cell tracking across biological scales. Nature Methods 2025 1–14 (2025) doi:10.1038/s41592-025-02778-0.

92. Young, G. & Coktas, A. Jex: High Performance Computing at the MRC Laboratory of Medical Sciences. 10.5281/ZENODO.10245365 doi: 10.5281/ZENODO.10245365.

93. Love, M. I., Huber, W. & Anders, S. Moderated estimation of fold change and dispersion for RNA-seq data with DESeq2. Genome Biol 15, 1–21 (2014).

94. Patro, R., Duggal, G., Love, M. I., Irizarry, R. A. & Kingsford, C. Salmon provides fast and bias-aware quantification of transcript expression. Nature Methods 2017 14:*4* **14**, 417–419 (2017).

95. Soneson, C., Love, M. I. & Robinson, M. D. Differential analyses for RNA-seq: transcript-level estimates improve gene-level inferences. F1000Res 4, 1521 (2016).

96. Love, M. I., Huber, W. & Anders, S. Moderated estimation of fold change and dispersion for RNA-seq data with DESeq2. Genome Biol 15, 1–21 (2014).

97. Ignatiadis, N., Klaus, B., Zaugg, J. B. & Huber, W. Data-driven hypothesis weighting increases detection power in genome-scale multiple testing. Nature Methods 2016 13:*7* **13**, 577–580 (2016).

98. Stephens, M. False discovery rates: a new deal. Biostatistics 18, 275–294 (2017).

99. McGuinness, B. E. et al. Regulation of APC/C Activity in Oocytes by a Bub1- Dependent Spindle Assembly Checkpoint. Current Biology 19, 369–380 (2009).

100. Madeira, F. et al. The EMBL-EBI Job Dispatcher sequence analysis tools framework in 2024. Nucleic Acids Res 52, W521–W525 (2024).

