## Extended data figures for "Wee1 opposes APC/C^Cdh1^ activity to promote S-phase entry"

Vuillemenot *et al.*

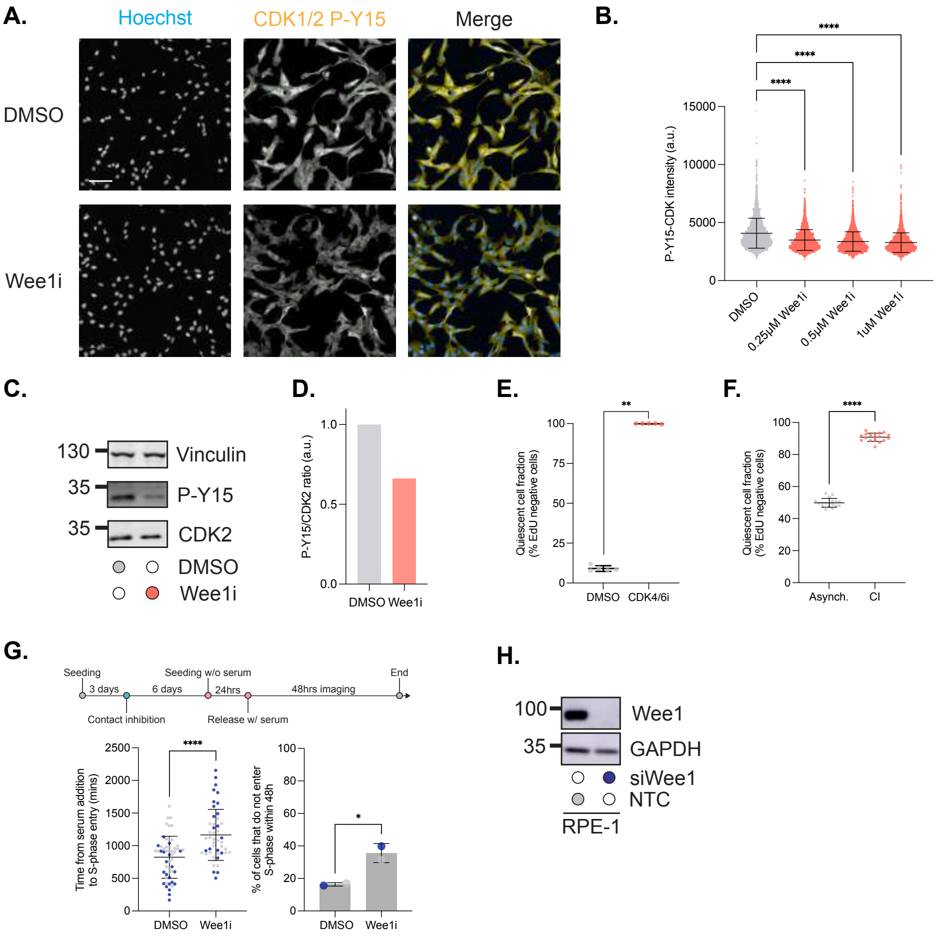


**Extended data Figure 1**

**Extended data Figure 1 (relating to Figure 1).**

**A.** Images of hTert-RPE1 cells treated with DMSO or 1 μM Wee1i for 1 hr. Hoechst is used to label nuclei (blue) and anti-CDK-pTyr15 (yellow) used to quantify the effect of Wee1 inhibition. Scale bar is 100 μm. **B.** Quantification of reduction in nuclear CDK-pTyr15 levels measured by immunostaining at different doses of Wee1i. One-way ANOVA test followed by Šídák's multiple comparisons test for significance, ****p<0.001. **C.** Western blot showing reduction in CDK-pTyr15 after 1 hr treatment with 1 μM Wee1i. Vinculin is used as a loading control. **D.** Quantification of western blot in (C) showing reduction in CDK-pTyr15 levels normalised to total CDK2 levels. **E.** Graph showing robust G0/G1 arrest (EdU negative cells) in hTert-RPE1 cells treated with CDK4/6i (Palbociclib) for 24 hr then pulsed with EdU for 24h. **F.** Graph showing robust G0/G1 arrest (EdU negative cells) in hTert-RPE1 cells grown to contact inhibition. EdU was pulsed in for the last 24 hr of contact inhibition to label actively proliferating cells. **G.** Graphs show length of G1 phase (left) and fraction of cells that do not enter S-phase (right) after Wee1i treatment upon release back into proliferation from contact inhibition followed by serum starvation. In E-G, mean +/- stdev are plotted with black lines, individual data points are shown, coloured by experimental repeat. Unpaired student's t-test for significance. ****p<0.0001; ***p<0.001; **p<0.01; *p<0.05. **H.** Western blot showing depletion of Wee1 protein by siRNA after 6 hr. GAPDH is used as a loading control.

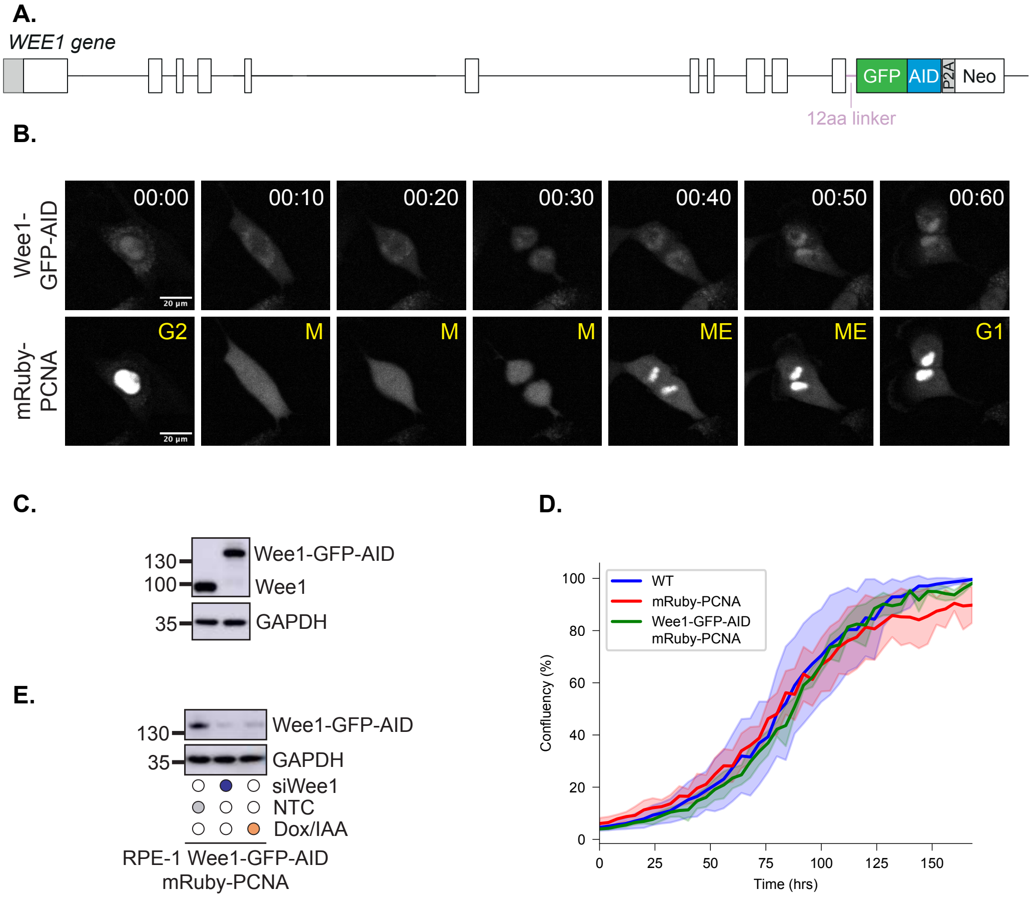


**Extended data Figure 2**

29

**Extended data Figure 2 (relating to Figure 1).**

**A.** Schematic for C-terminal tagging of the endogenous *WEE1* gene with GFP-AID tag. **B.** Images showing correct nuclear localisation of Wee1-GFP-AID, as previously described[1]. M is mitosis, ME is mitotic exit. **C.** Western blot showing homozygous tagging of endogenous Wee1. GAPDH is used as a loading control. **D.** Growth curve of parental (wild-type, WT, in blue), mRuby-PCNA only (in red) and Wee1-GFP-AID mRuby-PCNA cells (in green). Mean and 95% confidence interval around the mean is shown. **E.** Western blot showing degradation of Wee1 protein after addition of Dox/IAA for 1h, to a similar level as with Wee1 targeting siRNA. GAPDH is used as a loading control.

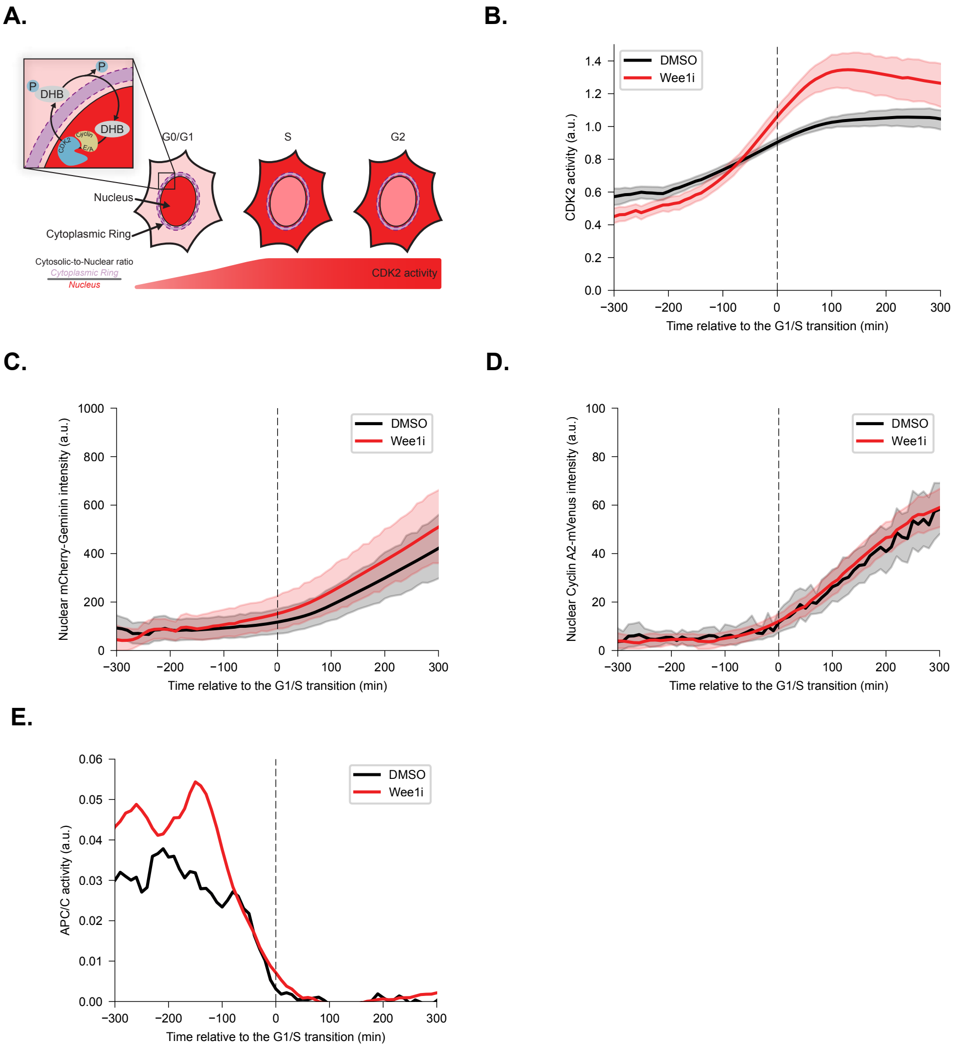


**Extended data Figure 3**

**Extended data Figure 3 (relating to Figure 2).**

**A.** Cartoon showing how the DHB-mCherry CDK2 activity sensor works. **B.** Graph shows CDK2 activity plotted relative to the G1/S transition for cells released from G0/G1 arrest in the presence of DMSO (control, black line) or Wee1i (red line). Mean and 95% confidence interval around the mean over time is shown. **C.** Graph shows timing and rate of mCherry-Geminin expression, plotted relative to the G1/S transition for cells released from G0/G1 arrest in the presence of DMSO (control, black line) or Wee1i (red line). Mean and 95% confidence interval around the mean over time is shown. These data were used to calculate $APC/C^{Cdh1}$ inactivation in Figure 2C. **D.** Graph shows timing and rate of CyclinA2-mVenus expression, plotted relative to the G1/S transition for cells released from G0/G1 arrest in the presence of DMSO (control, black line) or Wee1i (red line). Mean and 95% confidence interval around the mean over time is shown. These data were used to calculate $APC/C^{Cdh1}$ inactivation in Figure S3E. **E.** Graph shows timing and rate of $APC/C^{Cdh1}$ inactivation based on a CyclinA2-mVenus readout, plotted relative to the G1/S transition for cells released from G0/G1 arrest in the presence of DMSO (control, black line) or Wee1i (red line). Mean over time is shown.

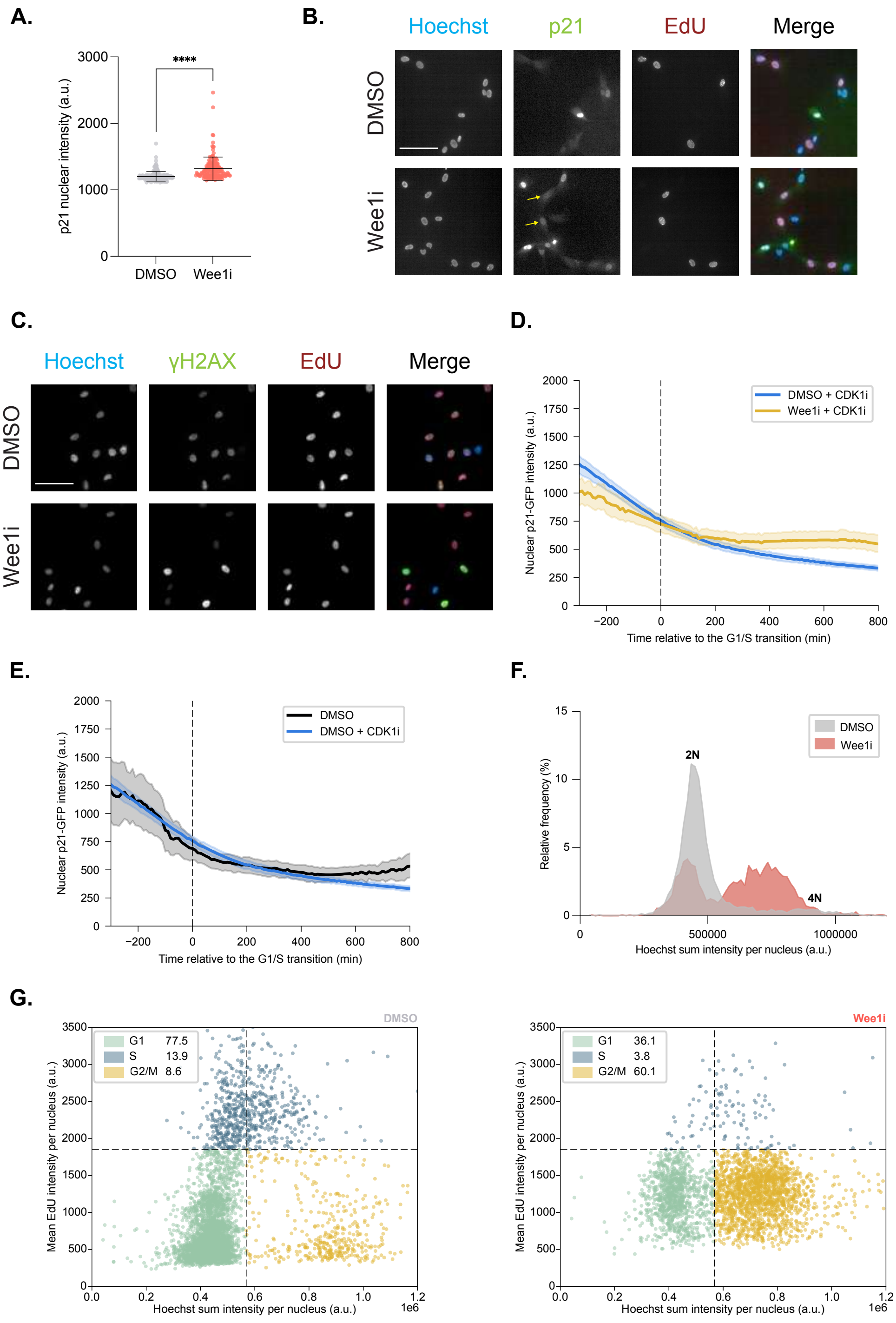


**Extended data Figure 4**

58

**Extended data Figure 4 (relating to Figure 3).**

**A.** Graph shows quantification of p21 intensity in S-phase nuclei from cells fixed 24 hr after release from CDK4/6i and stained for EdU (to identify S-phase cells) and p21. Unpaired student's t-test for significance. ****p<0.0001. **B.** Images show p21 immunostaining and EdU positivity in cells fixed 24h after release from CDK4/6i. Hoechst is in blue, p21 is in green and EdU is in red in merged images. Yellow arrows indicate cells in S-phase (EdU positive) with p21 present in the nucleus. **C.** Images show $\gamma$H2AX immunostaining and EdU positivity in cells fixed 24h after release from CDK4/6i. Hoechst is in blue, $\gamma$H2AX is in green and EdU is in red in merged images. Scale bars are 100 $\mu$m. **D.** Quantification of p21-GFP nuclear intensity over time, plotted relative to the G1/S transition for cells released from G0/G1 arrest in the presence of CDK1i (blue line) or combined Wee1i and CDK1i (yellow line). Mean and 95% confidence interval around the mean over time is shown. **E.** Quantification of p21-GFP nuclear intensity over time, plotted relative to the G1/S transition for cells released from G0/G1 arrest in the presence of DMSO (black line) or CDK1i (blue line). Mean and 95% confidence interval around the mean over time is shown. **F.** Hoechst sum intensity plot of cells showing cell cycle profile in cells 24h after released from CDK4/6i. Wee1i-treated cells do not completely replicate their DNA and accumulate between 2n and 4n populations. **G.** Hoechst sum intensity plotted against mean EdU intensity per nucleus. DMSO treated cells in the upper panel, Wee1i treated cells in the lower panel. EdU was pulsed in for the last 30 mins before fixation. Wee1i treated cells between the 2n and 4n populations are not actively replicating their DNA and have exited S-phase with incompletely replicated DNA.

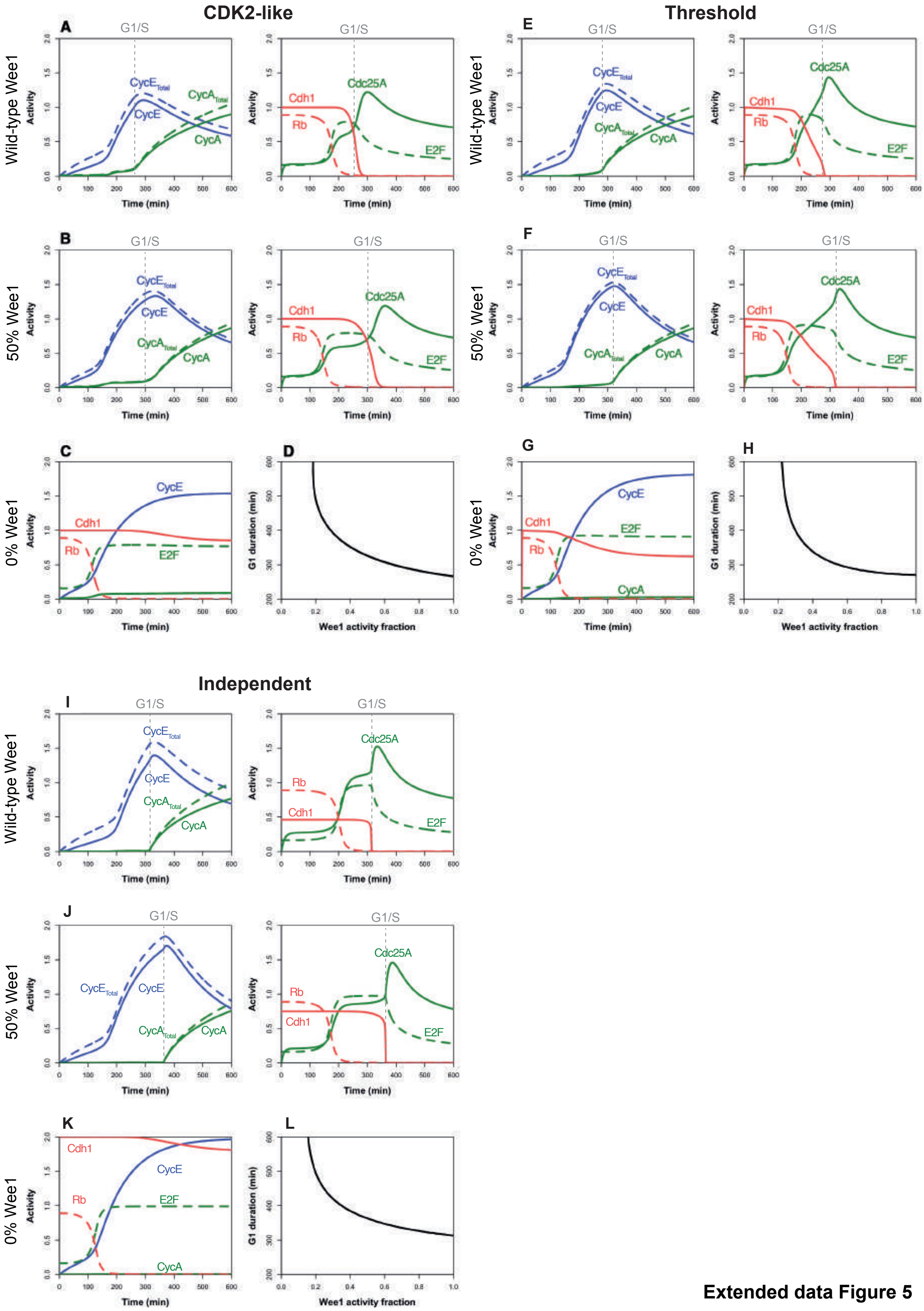


Extended data Figure 5

82

**Extended data Figure 5 (relating to Figure 4).**

Output from “CDK2-like” ODE model for Wee1 inhibition of Cdh1 for wild-type Wee1 activity (**A**), 50% of Wee1 activity (**B**) and 0% Wee1 activity (**C**). **D.** G1 length plotted as a function of Wee1 activity showing an infinite G1 length (G0 arrest) at very low levels of Wee1 activity. Output from “threshold” ODE model for Wee1 inhibition of Cdh1 for wild-type Wee1 activity (**E**), 50% of Wee1 activity (**F**) and 0% Wee1 activity (**G**). **H.** G1 length plotted as a function of Wee1 activity showing an infinite G1 length (G0 arrest) at very low levels of Wee1 activity. Output from “independent” ODE model for Wee1 inhibition of Cdh1 for wild-type Wee1 activity (**I**), 50% of Wee1 activity (**J**) and 0% Wee1 activity (**K**). **L.** G1 length plotted as a function of Wee1 activity showing an infinite G1 length (G0 arrest) at very low levels of Wee1 activity.

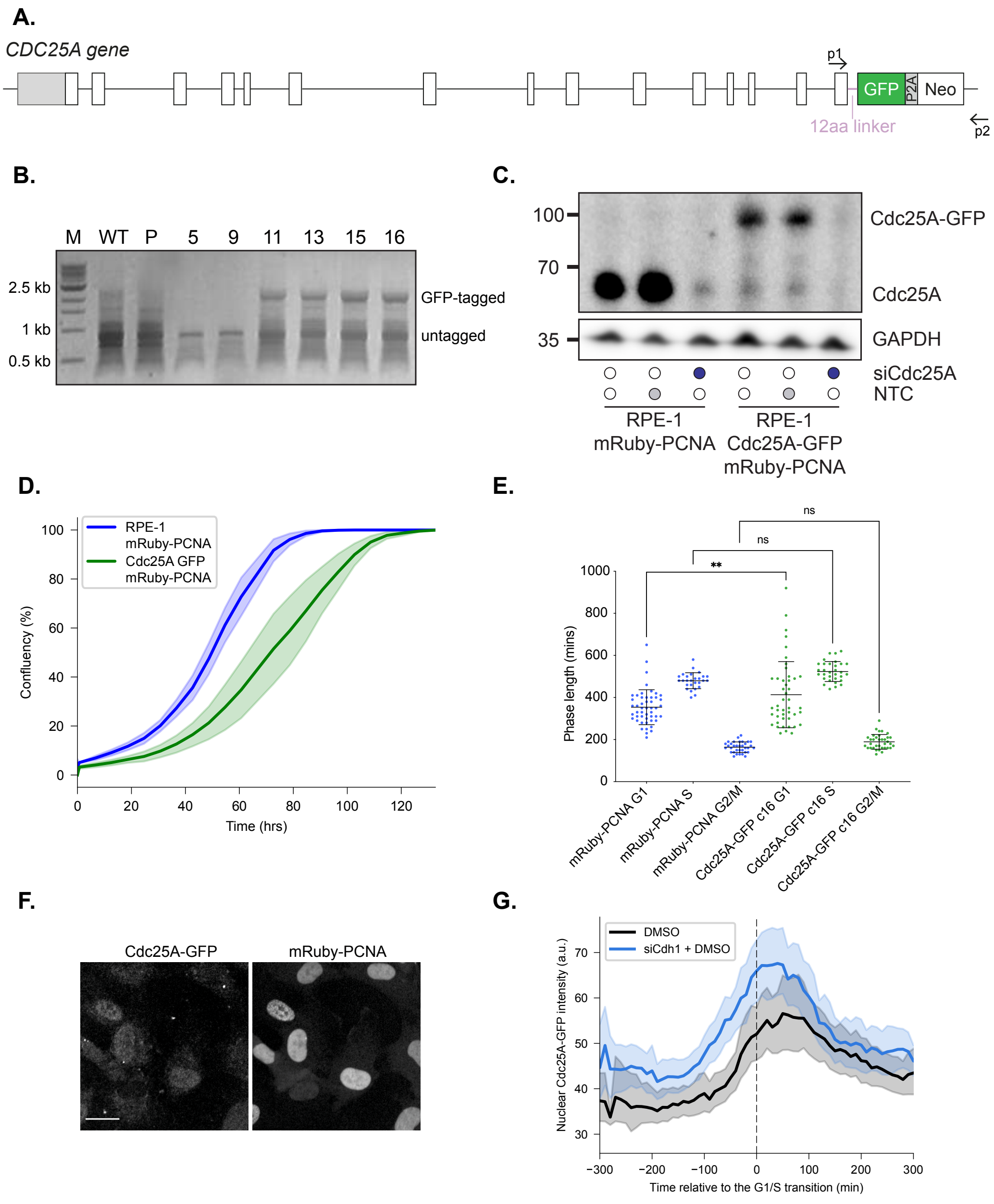


**Extended data Figure 6**

**Extended data Figure 6 (relating to Figure 4).**

**A.** Schematic for C-terminal tagging of the *CDC25A* gene with a GFP tag. **B.** Genomic DNA PCR with primers indicated in A (p1 and p2) demonstrating heterozygous tagging of Cdc25A with GFP. WT refers to wild-type hTert-RPE1. P refers to the parental hTert-RPE1 mRuby-PCNA cells. Numbers refer to different clones. Clone 16 was used for the experiments shown. **C.** Western blot showing specific heterozygous tagging of Cdc25A protein with GFP. GAPDH is used as a loading control. **D.** Growth curves of parental mRuby-PCNA only (in blue) and Cdc25A-GFP mRuby-PCNA cells (in green). Mean and 95% confidence interval around the mean is shown. **E.** Graph shows cell cycle phase timing (G1, S and G2/M) between parental mRuby-PCNA only cells and Cdc25A-GFP mRuby-PCNA cells. One-way ANOVA for significance. **$p<0.01$, ns is not significant. **F.** Images showing correct nuclear localisation of Cdc25A-GFP. Scale bar is 10 μm. **G.** Quantification of Cdc25A-GFP nuclear intensity over time, plotted relative to the G1/S transition for cells transfected with non-targeting control (NTC, black line) or Cdh1-targeting (blue line) siRNA. released from G0/G1 arrest in the presence of DMSO (control, black line) or Wee1i (red line). Mean and 95% confidence interval around the mean over time is shown.

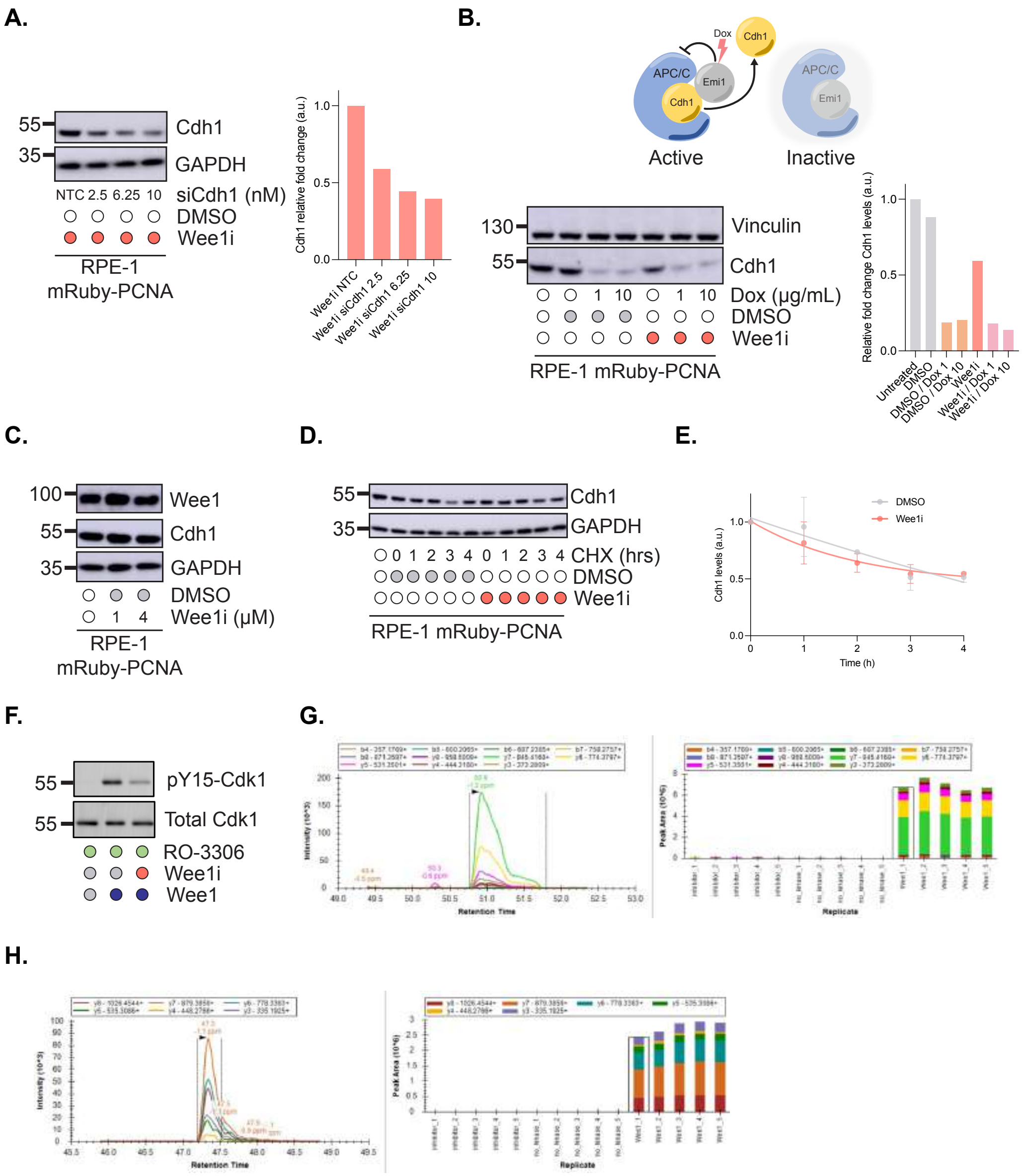


Extended data Figure 7

**Extended data Figure 7 (relating to Figure 4).**

**A.** Western blot showing Cdh1 depletion by siRNA 6h after transfection with different concentrations of siRNA. GAPDH is used as a loading control. Graph shows quantification of Cdh1 depletion normalised to GAPDH and relative to the NTC condition. **B.** Top: cartoon shows how doxycycline-induced overexpression of Emi1 promotes Cdh1 dissociation from APC/C and subsequent degradation. Middle: western blot showing decrease in Cdh1 protein after induction of Emi1 expression with Dox for 4 hr. Vinculin is used as a loading control. Right: quantification of Cdh1 depletion normalised to vinculin and relative to the untreated condition. **C.** Western blot showing Cdh1 protein levels in cells treated with 1 or 4 μM of Wee1i for 6h. GAPDH is used as a loading control. **D.** Western blot showing stability of Cdh1 protein in G0/G1 arrested cells after Wee1i treatment. GAPDH is used as a loading control. Cells were held in G0/G1 in order to compare Cdh1 protein stability at the same cell cycle state in DMSO versus Wee1i treated cells. **E.** Graph plotting Cdh1 protein levels from (D) with a one-phase decay in DMSO- versus Wee1i-treated cells. **F.** Westen blot showing positive control reaction for the *in vitro* kinase assay for Wee1 and CyclinB/CDK1. An antibody raised against CDK phospho-tyrosine 15 (pY15-Cdk1) was used to probe for phosphorylation of CDK1 by Wee1. CDK1 inhibitor (RO-3306) was added to prevent CDK1-mediated inhibition of Wee1 kinase. **G.** Validation of Y91 site by targeted mass spectrometry of DGLAY[Phospho (Y)]SALLK 2+ peptide. Note presence of y6 fragment that confirms Y phosphorylation position **H.** Validation of Y130 site by targeted mass spectrometry of GLFTY[Phospho (Y)]SLSTK 2+ peptide. Note presence of y6 fragment that confirms Y phosphorylation position.

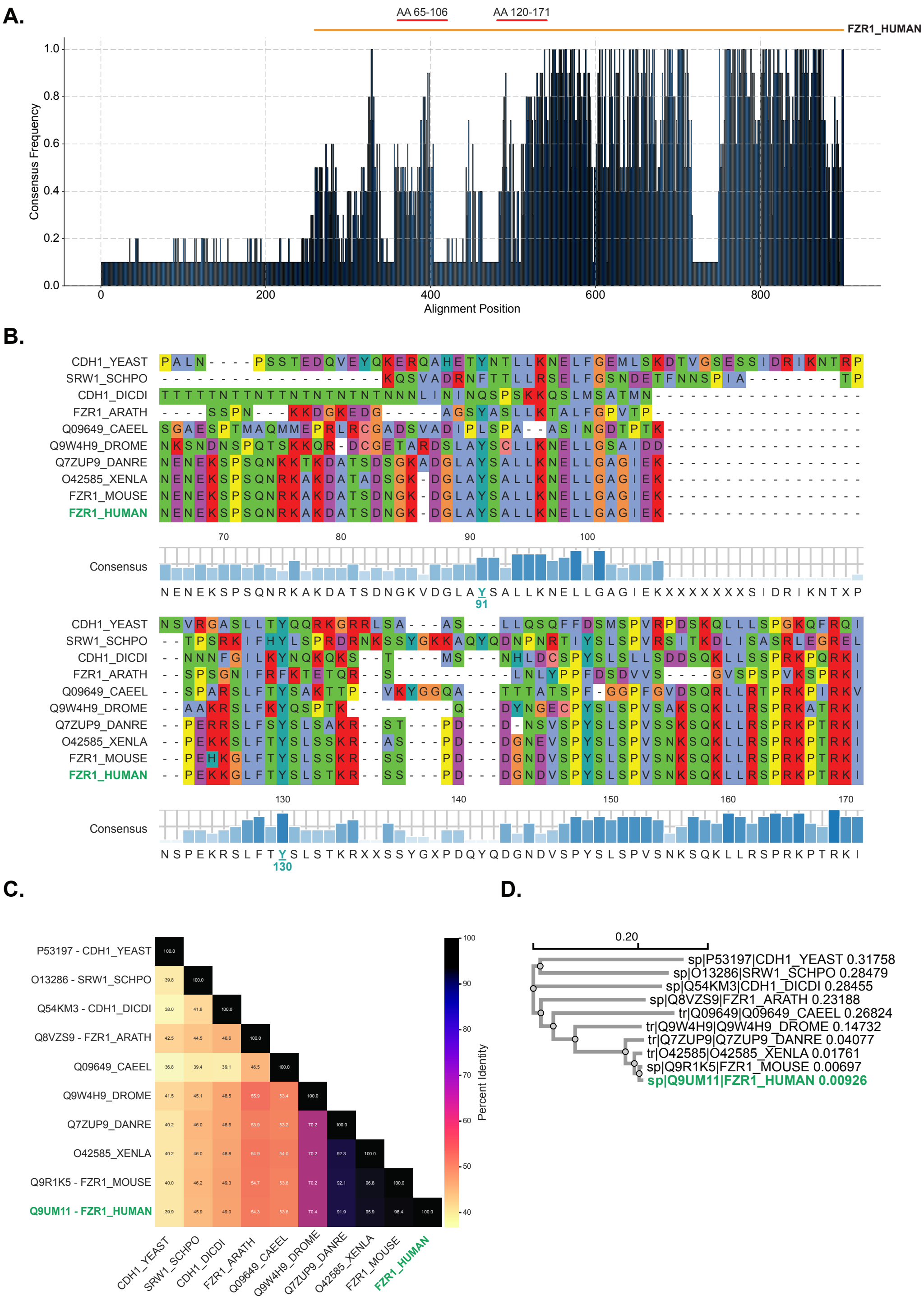


**Extended data Figure 8**

**Extended data Figure 8 (relating to Figure 4).**

**A.** Consensus frequency of Cdh1 residues across species aligned with Clustal Omega. Orange line above the chart indicates position of human Cdh1 (encoded by *FZR1*). Red lines above chart indicate portions of the alignment shown in (B). **B.** Portions of Clustal Omega alignment shown in (A) of Cdh1 across species with consensus frequency. Top: amino acids 65-106 to show conservation of Y91. Bottom: amino acids 120-171 to show conservation of Y130. **C.** Cdh1 identity matrix. **D.** Cdh1 phylogenetic tree.

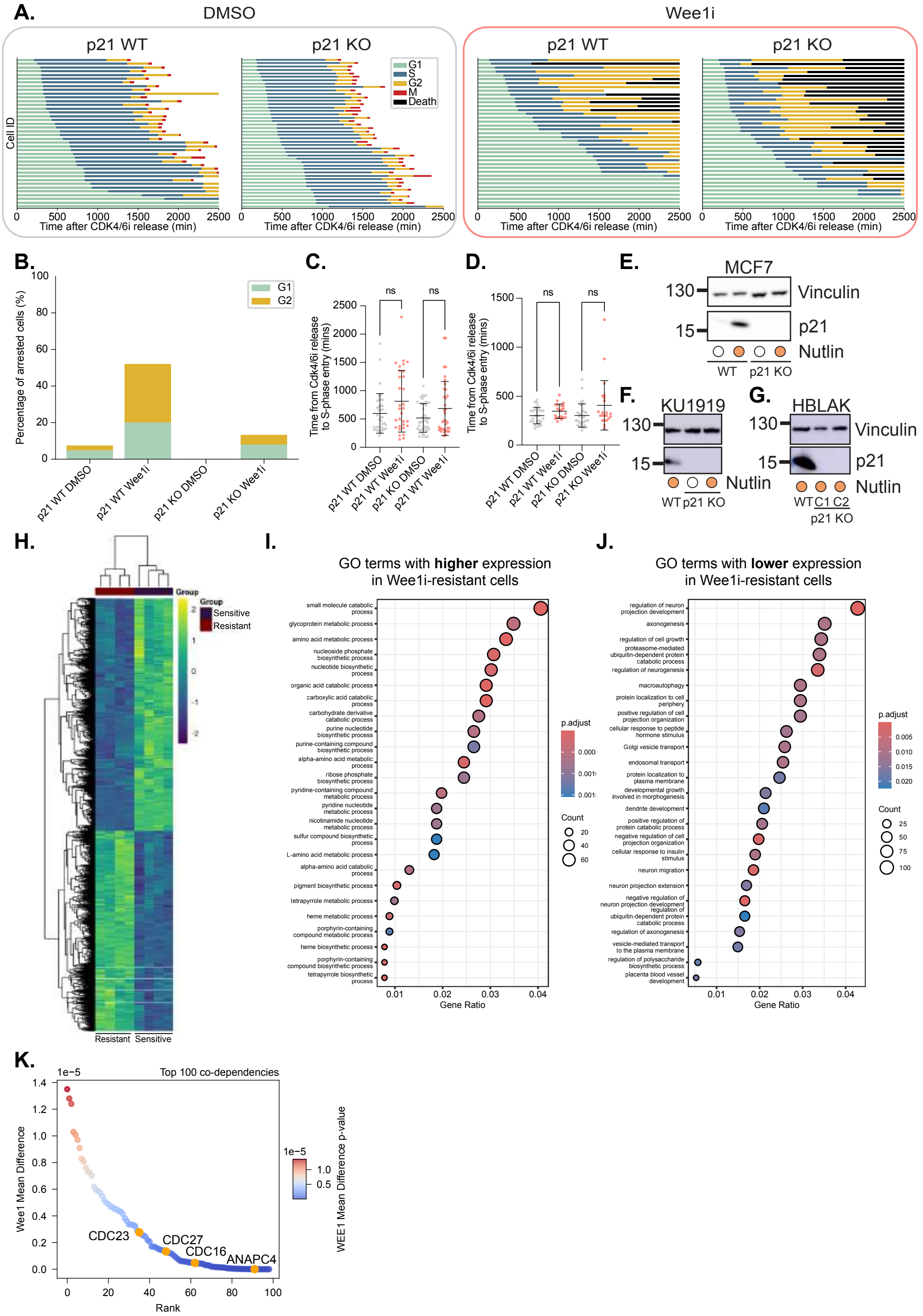


**Extended data Figure 9**

**Extended data Figure 9 (relating to Figure 5).**

**A.** Graphs show cell cycle phase timings for p21 wild-type (WT) and p21 knockout (KO) MCF7 cells treated with DMSO (left) or Wee1i (right) after release from CDK4/6i. **B.** Graph shows fraction of MCF7 cells arresting in G1 or G2 phase in p21WT and p21KO cells treated with DMSO or Wee1i from cells plotted in (A). **C.** Graphs show length of G1 phase (left) for p21 wild-type (WT) and p21 knockout (KO) MCF7 cells after Wee1i treatment upon release from CDK4/6i-mediated G0/G1 arrest. One-way ANOVA followed by Tukey's multiple comparisons test for significance. **D.** Graphs show length of G1 phase (left) for p21 wild-type (WT) and p21 knockout (KO) KU1919 cells after Wee1i treatment upon release from CDK4/6i-mediated G0/G1 arrest. One-way ANOVA followed by Tukey's multiple comparisons test for significance. **E-G.** Western blots showing the validation of p21 knock-out (KO) in MCF7 (E), KU1919 (F) and HBLAK (G) cell lines. Vinculin is used as a loading control. **H.** Heatmap showing significantly differentially expressed genes between Wee1i resistant (left) and Wee1i sensitive (right) cells. Four technical repeats were performed for each condition. **I.** Pathway analysis of upregulated genes in Wee1i-resistant cells compared to parental cells. **J.** Pathway analysis of downregulated genes in Wee1i-resistant cells compared to parental cells. **K.** Graph shows data replotted from DepMap of the top 100 genes that synergise with Wee1 loss. APC/C components are labelled. Four of the 19 APC/C subunits are in the list.

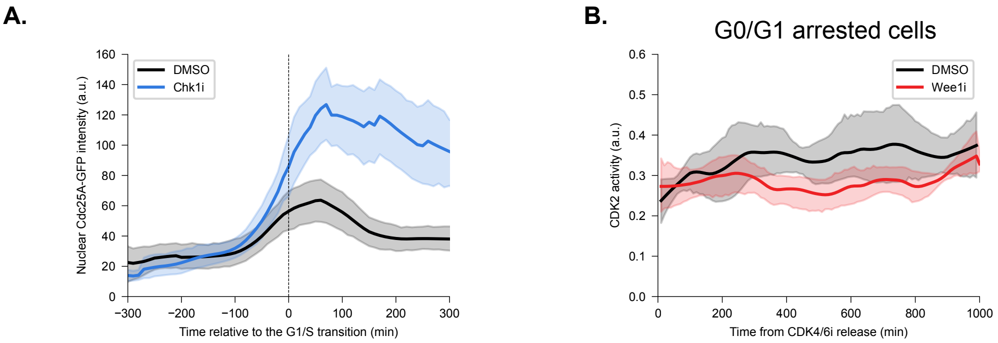


**Extended data Figure 10**

166

**Extended data Figure 10 (relating to Discussion).**

**A.** Graph shows nuclear Cdc25A-GFP intensity calculated from hTert-RPE1 mRuby-PCNA Cdc25A-GFP cells released from contact-inhibited and serum-starved G0 cells plotted around the G1/S transition. Upon release, cells were treated with DMSO or 1 μm of the Chk1i, LY2880070, before timelapse imaging was started. Mean and 95% confidence interval around the mean over time is shown. **B.** Graph shows CDK2 activity as measured by the CDK2L-GFP sensor in cells that remain arrested in G0/G1 after DMSO or Wee1i treatment. Data taken from same experiment as Figure 2B. Mean and 95% confidence interval around the mean over time is shown.
