## Supplementary Material for "Wee1 opposes APC/C^Cdh1^ activity to promote S-phase entry"

### Supplementary Movies

**Supplementary Movie 1.** hTert-RPE1 mRuby-PCNA (magenta) CDK2L-GFP (gold) expressing cells released from CDK4/6i arrest into DMSO (left) or Wee1i (right). In cells treated with Wee1i, CDK2 activity starts to increase earlier and to a higher level during G1 than in DMSO-treated cells.

**Supplementary Movie 2.** hTert-RPE1 mRuby-PCNA (red) p21-GFP (green) expressing cells released from CDK4/6i arrest into DMSO (left) or Wee1i (right). In cells treated with Wee1i, p21-GFP starts to accumulate in S-phase nuclei when mRuby-PCNA foci are still present.

**Supplementary Movie 3.** hTert-RPE1 mRuby-PCNA (magenta) Cdc25A-GFP (gold) cells released from CDK4/6i. A pulse of nuclear Cdc25A-GFP expression is visible around the G1-S transition.

### Supplementary tables

Supplementary table 1 – Log gene expression in MCF WT (wild-type, Wee1i sensitive) and MCF7 MK (Wee1i resistant).

Supplementary table 2 – Cell lines

| Cell line | Construct | Species | Tissue | Type | Media | Antibio-tics | Cat no. | Source |
|---|---|---|---|---|---|---|---|---|
| **hTert-RPE1** | - | Human | Epithelial (Retina) | Normal | DMEM | - | CRL-2302 | ATCC |
| **hTert-RPE1** | mRuby-PCNA | Human | Epithelial (Retina) | Normal | DMEM | - | - | Mansfeld lab[2] |
| **hTert-RPE1** | mRuby-PCNA CDK2L-GFP | Human | Epithelial (Retina) | Normal | DMEM | - | - | Barr lab[3] |
| **hTert-RPE1** | DHB-mCherry mTurq2-PCNA Cdc6-mVenus | Human | Epithelial (Retina) | Normal | DMEM | - | - | Cook lab[4] |
| **hTert-RPE1** | mRuby-PCNA Cdc25A-GFP | Human | Epithelial (Retina) | Normal | DMEM | - | - | Barr lab, this study |
| **hTert-RPE1** | myc-OsTIR1 | Human | Epithelial (Retina) | Normal | DMEM | - | - | Hochegger lab[5] |
| **hTert-RPE1** | myc-OsTIR1 Wee1-GFP-AID mRuby-PCNA | Human | Epithelial (Retina) | Normal | DMEM | - | - | Barr lab, this study |
| **hTert-RPE1** | PIP-Fucci Cdt1-geminin mTurq2-PCNA | Human | Epithelial (Retina) | Normal | DMEM | - | - | Cook lab[6] |
| **hTert-RPE1** | p21-GFP mRuby-PCNA | Human | Epithelial (Retina) | Normal | DMEM | - | - | Barr lab[7] |
| **hTert-RPE1** | CCNB1-eYFP mTurq2-EMI1 53BP1-mCherry | Human | Epithelial (Retina) | Normal | DMEM | - | - | Medema lab[8] |
| **hTert-RPE1** | CCNA2-mVenus mRuby-PCNA | Human | Epithelial (Retina) | Normal | DMEM | - | - | Pines lab[9] |
| **MCF7** | - | Human | Epithelial (Breast) | Cancer | DMEM | - | HTB-22 | ATCC |

| | | | | | | | | |
|---|---|---|---|---|---|---|---|---|
| **MCF7** | mRuby-PCNA | Human | Epithelial (Breast) | Cancer | DMEM | - | - | Barr lab, this study |
| **MCF7** | mRuby-PCNA p21 KO | Human | Epithelial (Breast) | Cancer | DMEM | - | - | Barr lab, this study |
| **KU1919** | - | Human | Epithelial-like (Bladder) | Cancer | RPMI | - | ACC 395 | DSMZ |
| **KU1919** | mRuby-PCNA p21-GFP | Human | Epithelial-like (Bladder) | Cancer | RPMI | - | - | Barr lab, this study |
| **KU1919** | mRuby-PCNA p21 KO | Human | Epithelial-like (Bladder) | Cancer | RPMI | - | - | Barr lab, this study |
| **HBLAK** | - | Human | Epithelial (Bladder) | Normal | Cnt Prime | - | - | CELLnTEC |
| **HBLAK** | mRuby-PCNA p21 KO | Human | Epithelial (Bladder) | Normal | Cnt Prime | - | | Barr lab, this study |

Supplementary table 3 - Plasmids

| Plasmid | Backbone | Insert | Source |
|---|---|---|---|
| **pX459 Wee1 CT gRNA** | pX459 pSpCas9[10] | sgRNA Wee1 | Barr Lab, this study |
| **pSK Wee1-GFP-AID-P2A-Neo** | pSKII- | Wee1-GFP-AID-P2A-Neo | Barr Lab, this study |
| **pX459 Cdc25A CT gRNA** | pX459 pSpCas9[10] | sgRNA Cdc25A | Barr Lab, this study |
| **pSK Cdc25A-GFP-Neo** | pSKII- | Cdc25A-GFP-P2A-Neo | Barr Lab, this study |

Supplementary table 4 - Drugs

| Drug | Target | Final concentration | Solvent | Cat no. | Source |
|---|---|---|---|---|---|
| **MK-1775** | Wee1 | 1 µM | DMSO | S1525 | Selleckchem |
| **Palbociclib** | CDK4/6 | 0.5-1 µM | DMSO | S4482 | Selleckchem |
| **RO-3306** | CDK1 | 7.5-18 µM | DMSO | S7747 | Selleckchem |
| **Doxycycline** | - | 1 µg.mL$^{-1}$ | ddH2O | D9891 | Sigma Aldrich |
| **IAA** | - | 500 µM | ddH2O | 87-51-4 | Sigma Aldrich |
| **EdU** | - | 5-10 µM | ddH2O | A10044 | ThermoFisher Scientific |
| **G418** | - | 1 mg/mL | ddH2O | 10131027 | ThermoFisher Scientific |

Supplementary table 5 - siRNA

| Target | Final concentration | Cat no. | Source |
|---|---|---|---|
| **NTC** | 20 nM | D-001830-01 | Horizon discovery |
| **FZR1 (Cdh1)** | 10 nM | 603619 | Horizon discovery |
| **WEE1 (Wee1)** | 20 nM | 16708 | Horizon discovery |
| **CDC25A (Cdc25A)** | 20 nM | 4390771 | Horizon discovery |

Supplementary table 6 - Antibodies

| Antibody | Species | Sources | Cat. | Dilution WB | Dilution IF |
|---|---|---|---|---|---|
| **Phospho-γH2AX (S139)** | Rabbit | Cell Signaling Technology | 2577 | | 1:1000 |
| **Y15-pCDK1/2/3/5** | Rabbit | Abcam | ab76146 | 1:1000 | 1:200 |
| **CDK1** | Mouse | Cell Signaling Technology | 9116 | 1:1000 | |
| **CDK2** | Mouse | Santa Cruz Biotechnology | Sc6248 | 1:200 | 1:500 |
| **pTyr-100** | Mouse | Cell Signaling Technology | 9411 | 1:2000 | |
| **p21** | Mouse | BD Biosciences | 556430 | 1:500 | 1:500 |
| **GAPDH** | Mouse | Cell Signaling Technology | 97166 | 1:1000 | |
| **Fzr1 (Cdh1)** | Mouse | Santa Cruz Biotechnology | sc56312 | 1:200 | |
| **Vinculin** | Rabbit | Cell Signaling Technology | 13901 | 1:1000 | |
| **Wee1** | Rabbit | Cell Signaling Technology | 4936 | 1:250 | |
| **MCM7** | Mouse | Santa Cruz Biotechnology | Sc56324 | | 1:500 |
| **Anti-rabbit HRP-linked** | Goat | Cell Signaling Technology | 7074P2 | 1:1000 | |
| **Anti-mouse HRP-linked** | Goat | Cell Signaling Technology | 7076P2 | 1:1000 | |
| **Anti-rabbit Alexa Fluor-488** | Goat | Invitrogen | A11008 | 1:1000 | 1:1000 |
| **Anti-mouse Alexa Fluor-488** | Goat | Invitrogen | A11001 | | 1:1000 |
| **Anti-rabbit Alexa Fluor-568** | Goat | Invitrogen | A11011 | | 1:1000 |
| **Anti-mouse Alexa Fluor-568** | Goat | Invitrogen | A11004 | | 1:1000 |
| **Anti-rabbit Alexa Fluor-647** | Goat | Invitrogen | A21245 | | 1:1000 |

| | | | | | |
|---|---|---|---|---|---|
| **Anti-mouse Alexa Fluor-647** | Goat | Invitrogen | A21235 | 1:1000 | 1:1000 |

Supplementary table 7 – Purified proteins

| Target | Cat no. | Source |
|---|---|---|
| **GST-Wee1** | PV3817 | ThermoFisher Scientific |
| **GST-CDK1+GST-CyclinB1** | ab271456 | Abcam |
| **Strep-Strep-Cdh1** | - | Alfieri lab[11] |

### Supplementary methods

*Modelling and parameters*

A base model of the G1/S transition

Our cell cycle model consists of coupled ODEs describing the kinetics of regulators relevant for the G1/S transition. A number of species were assumed to be in pseudo-steady state if they engage in 'fast' reactions relative to protein synthesis and degradation. These species were modelled as algebraic functions of the other regulators.

We begin by describing the synthesis and degradation of E2F-regulated genes. The Cdc25A phosphatase is synthesised basally, as well as in a manner that is stimulated by the transcription factor. Similarly, the degradation of the protein takes place basally and under the action of the ubiquitin ligase APC/C:Cdh1

$$\frac{dCdc25T}{dt} = ks25' + ks25" \cdot E2F - (kd25' + kd25" \cdot Cdh1) \cdot Cdc25T \qquad (1)$$

Where $ks25'$, $ks25"$, $kd25'$ and $kd25"$ and kinetic constants. The E2F transcription factor also stimulates the production of Cyclin E and Cyclin A. Importantly, only Cyclin A is a Cdh1 substrate, such that we model the two cyclins as follows:

$$\frac{dCycET}{dt} = kse' + kse \cdot \mathrm{E2F} - \mathrm{kde'} \cdot (\mathrm{CycET} - \mathrm{CycE}) - \mathrm{kde} \cdot CycE \qquad (2)$$

$$\frac{dCycAT}{dt} = ksa * E2F - (kda' + kda \cdot Cdh1) \cdot CycAT \qquad (3)$$

Notice that the active, dephosphorylated forms of the two cyclins are referred to as CycE and CycA, respectively, in opposition to the total concentrations (CycET and CycAT). Furthermore, we assume that dephosphorylated form of CycE is degraded at a faster rate[12]. The phosphorylation and dephosphorylation of these cyclins is catalysed by Wee1 and Cdc25, as indicated by the ODEs:

$$\frac{dCycE}{dt} = kse' + kse \cdot E2F - kwee \cdot CycE + V25 \cdot (CycET - CycE) - kde \cdot CycE \quad (4)$$

$$\frac{dCycA}{dt} = ksa * E2F - kwee \cdot CycA + V25 \cdot (CycAT - CycA) - (kda' + kda \cdot Cdh1) \cdot CycA \quad (5)$$

As we assume that Wee1 is not regulated before the G2/M transition, its activity is given by a constant *kwee*, nominally equal to 1, unless otherwise stated. *V25* represents rate function of Cdc25, which accounts for the activity of CDK phosphorylated and dephosphorylated forms:

$$V25 = k25' \cdot (Cdc25T - Cdc25P) + k25'' \cdot Cdc25P \quad (6)$$

Where the phosphorylated Cdc25 is given by:

$$Cdc25P = Cdc25T \frac{CycE^p}{J25^p + CycE^p} \quad (7)$$

The steady state phosphorylation of Cdc25 is modelled by a CycE-dependent Hill function. The nonlinearity emerges thanks to distributive multisite phosphorylation by the kinase[13]. For the same reason, the G1/S transition inhibitors APC/C:Cdh1 ubiquitin ligase and the transcriptional repressor Rb are also modelled by Hill functions:

$$Cdh1 = Cdh1T \cdot \frac{\alpha^n}{\alpha^n + (CycE + \gamma \cdot CycA)^n} \quad (8)$$

$$Rbt = Rbtot \cdot \frac{\beta^m}{\beta^m \cdot (1 + SK) + (CycE + CycA)^m} \quad (9)$$

Where *Cdh1T* and *Rbtot* are the total protein concentrations, which are assumed to be constant. $\gamma$ is a parameter that accounts for the greater strength of Cdh1 inhibition by CycA. *SK* stands for 'Starter kinase', which corresponds to the activity of CycD:CDK4/6. Here, it is taken as a constant, nominally set to 1.25, corresponding to the mitogen stimulated state. Lowering this value simulates mitogen withdrawal, arresting the system in the G0 state, where there is negligible cyclin expression.

*Rbt*, the dephosphorylated Rb fraction, forms an inhibitory stoichiometric complex with the E2F transcription factor, as described by the steady state algebraic expression[14]:

$$Comp = \frac{BB2 - \sqrt{BB2^2 - 4 * Rbt * E2FT}}{2} \tag{10}$$

Where E2FT is the total concentration of E2F, taken as a constant and $BB2 = Rbt + E2FT + Kdiss$, where Kdiss is the dissociation constant between E2F and Rb.

E2F is also inhibited by CycA:CDK2-dependent phosphorylation in S-phase[15]. We model the steady state phosphorylated fraction of E2F as:

$$E2FPt = E2FT \cdot \frac{CycA}{kdpa + CycA} \tag{11}$$

As we assume Rb binding and CycA-dependent phosphorylation are independent, we express the 'active' E2F as:

$$E2F = \frac{(E2FT - E2FPt) \cdot (E2FT - Comp2)}{E2FT} \tag{12}$$

<u>The Cdk2-like inhibition model</u>

The Cdk2-like inhibition model assumes that Wee1 phosphorylates the same Cdh1 residues as CycA and CycE. Consequently, the Cdh1 activity function is updated so that its Hill function takes a weighted sum of the activities of CycE, CycA and Wee1 as input:

$$Cdh1 = Cdh1T \cdot \frac{\alpha^n}{\alpha^n + (CycE + \gamma \cdot CycA + kw * kwee)^n} \tag{8.1}$$

Note that the *kwee* parameter refers to the activity of Wee1 and *kw* is a dimensionless constant that represents the strength of Wee1 inhibition on Cdh1 relative to CycE.

The threshold model

This iteration of the model is based on the idea that CycE:Cdk2 to has little affinity for Cdh1, and therefore negligible activity towards it[16–18]. We hypothesised that the role of Wee1 might be to mediate the interaction between CycE and APC/C:Cdh1. Effectively, we assumed that Wee1 decreases the dissociation constat between the two proteins. Therefore, we decided to model Cdh1 as a differential equation, where APC/C: Cdh1 exists as a dephosphorylated form (Cdh1) and a phosphorylated form (Cdh1T – Cdh1). Both the forward and the backwards reactions are modelled as Michaelis-Menten processes[19], to account for the sharpness of the response to CDK2:

$$\begin{aligned} \frac{dCdh1}{dt} = kdp \frac{Cdh1T - Cdh1}{jp + (Cdh1T - Cdh1)} - kpa \cdot CycA \frac{Cdh1}{jpa + Cdh1} \\ - kpe \cdot CycE \frac{Cdh1}{jpe * \frac{kw}{kw + kwee} + Cdh1} \end{aligned} \tag{8.2}$$

The forward reaction (i.e. the dephosphorylation of Cdh1), which produces the active form, is catalysed by a constitutive phosphatase, whose activity is included in the rate constant *kdp*. The Michaelis constant of this reaction is *jp*. The phosphorylation reaction is driven by CycA with catalytic rate constant *kpa* and Michaelis constant *jpa*. The effect of CycE is modelled analogously, but notice that the Michaelis constant (the affinity between CycE and Cdh1) was written as a saturable process dependent on kwee, where the intrinsic Kd between CycE and Cdh1 decreases with Wee1 levels. This approach, though *ad hoc*, is completely analogous to the modelling of competitive inhibition, except Wee1 reduces rather than increases the apparent Michaelis constant.

Note that in this case, we have done away with the steady state assumption for Wee1, as the closed form algebraic expression was too unwieldy.

### The independent inhibition model

In this version, we assumed that Wee1 phosphorylates a separate subset of Cdh1 residues from Cdk2. Therefore, Wee1 would inactivate APC/C:Cdh1 via a CDK2-independent route. Formally, we expressed this as:

$$Cdh1 = Cdh1T \cdot \frac{\alpha^n}{\alpha^n + (CycE + \gamma \cdot CycA)^n} \cdot \frac{kw}{kw + kwee} \tag{8.3}$$

Essentially, we multiplied the Cdh1 expression in the base model by a factor that decreases hyperbolically with the activity of Wee1, *kwee*.

### Parameters

Supplementary table 8 – Numerical values of parameters in the base model

| Parameter | Description | Value |
|---|---|---|
| **kse'** | Basal synthesis rate of CycE | 0 |
| **kse"** | E2F-induced synthesis rate of CycE | 0.02 |
| **kde'** | Degradation rate of Wee1-phosphorylated CycE | 0.001 |
| **kde"** | Degradation rate of dephosphorylated CycE | 0.01 |
| **ksa** | E2F-induced synthesis rate of CycA | 0.01 |
| **kda'** | Basal degradation rate of CycA | 0.001 |
| **kda** | Cdh1-induced degradation rate of CycA | 0.1 |
| **kwee** | Wee1 activity | 1 |
| **k25'** | Activity of dephosphorylated Cdc25A | 0.1 |
| **k25"** | Activity of phosphorylated Cdc25A | 10 |
| **ks25'** | Basal synthesis rate of Cdc25A | 0.01 |
| **ks25"** | E2F-induced synthesis rate of Cdc25A | 0.1 |
| **kd25'** | Basal degradation rate of Cdc25A | 0.05 |
| **kd25"** | Cdh1-induced degradation rate of Cdc25A | 0.1 |
| **J25** | Threshold for Cdc25A activation by CycE | 0.1 |
| **p** | Hill exponent for CycE effect on Cdc25A | 1 |
| **Cdh1T** | Total Cdh1 concentration | 1 |
| **alpha** | Threshold for CycE/CycA inhibition of Cdh1 | 1.5 |
| **n** | Hill exponent of Cdh1 inhibition | 10 |
| **gamma** | Weighting factor for CycA contribution to Cdh1 inhibition | 4 |

| | | |
|---|---|---|
| **E2FT** | Total E2F concentration | 1 |
| **kdpa** | Threshold for E2F phosphorylation by CycA | 0.3 |
| **Rbtot** | Total Rb concentration | 2 |
| **beta** | Threshold for CycE/CycA inhibition of Rb | 0.3 |
| **m** | Hill exponent in Rb inhibition | 4 |
| **Kdiss** | Dissociation constant for Rb:E2F complex | 0.01 |
| **SK** | Starter Kinase (CycD:Cdk4/6) effect on Rb regulation | 1.25 |

The subsequent model variants had specific parameters modified as necessary to allow for a good qualitative fit of the experimental data. All parameter values are available in the supplementary .ode files.

Computation

The numerical integration and time course simulation of all models has been carried out in XPPAUT [20]. Estimates of G1/S timing over a range of Wee1 activities have been obtained by recording the time at which Cdh1 activity falls below the nominal value of 0.2, using the custom scripts Analysis.ode, available as supplementary files. All plots were generated by exporting data from regular and analysis .ode files, and processing using the R Statistical Software scripts[21] FigScript.R and SuppFigScrip.R.

### References


1. Baldin, V. & Ducommun, B. Subcellular localisation of human wee1 kinase is regulated during the cell cycle. *J Cell Sci* **108**, 2425–2432 (1995).

2. Zerjatke, T. *et al.* Quantitative Cell Cycle Analysis Based on an Endogenous All-in-One Reporter for Cell Tracking and Classification. *Cell Rep* **19**, 1953–1966 (2017).

3. Barr, A. R., Heldt, F. S., Zhang, T., Bakal, C. & Novák, B. A Dynamical Framework for the All-or-None G1/S Transition. *Cell Syst* **2**, (2016).

4. Matson, J. P. *et al.* Intrinsic checkpoint deficiency during cell cycle re-entry from quiescence. *Journal of Cell Biology* **218**, 2169–2184 (2019).

5. Hégarat, N. *et al.* Cyclin A triggers Mitosis either via the Greatwall kinase pathway or Cyclin B. *EMBO J* **39**, e104419 (2020).

6. Grant, G. D., Kedziora, K. M., Limas, J. C., Cook, J. G. & Purvis, J. E. Accurate delineation of cell cycle phase transitions in living cells with PIP-FUCCI. *Cell Cycle* **17**, 2496–2516 (2018).

7. Barr, A. R. *et al.* DNA damage during S-phase mediates the proliferation-quiescence decision in the subsequent G1 via p21 expression. *Nat Commun* **8**, (2017).

8. Hornsveld, M. *et al.* A FOXO-dependent replication checkpoint restricts proliferation of damaged cells. *Cell Rep* **34**, 108675 (2021).

9. Collin, P., Nashchekina, O., Walker, R. & Pines, J. The spindle assembly checkpoint works like a rheostat rather than a toggle switch. *Nat Cell Biol* **15**, 1378–1385 (2013).

10. Ran, F. A. *et al.* Genome engineering using the CRISPR-Cas9 system. *Nat Protoc* **8**, 2281 (2013).

11. Chang, L., Zhang, Z., Yang, J., McLaughlin, S. H. & Barford, D. Atomic structure of the APC/C and its mechanism of protein ubiquitination. *Nature 2015 522:7557* **522**, 450–454 (2015).

12. Won, K. A. & Reed, S. I. Activation of cyclin E/CDK2 is coupled to site-specific autophosphorylation and ubiquitin-dependent degradation of cyclin E. *EMBO J* **15**, 4182–4193 (1996).

13. Novak, B. & Tyson, J. J. Numerical analysis of a comprehensive model of M-phase control in Xenopus oocyte extracts and intact embryos. *J Cell Sci* **106**, 1153–1168 (1993).

14. Kim, J. K., Josić, K. & Bennett, M. R. The relationship between stochastic and deterministic quasi-steady state approximations. *BMC Syst Biol* **9**, 1–13 (2015).

15. Xu, M., Sheppard, K. A., Peng, C. Y., Yee, A. S. & Piwnica-Worms, H. Cyclin A/CDK2 binds directly to E2F-1 and inhibits the DNA-binding activity of E2F-1/DP-1 by phosphorylation. *Mol Cell Biol* **14**, 8420–8431 (1994).

16. Lukas, C. *et al.* Accumulation of cyclin B1 requires E2F and cyclin-A-dependent rearrangement of the anaphase-promoting complex. *Nature 1999 401:6755* **401**, 815–818 (1999).

17. Sørensen, C. S. *et al.* A Conserved Cyclin-Binding Domain Determines Functional Interplay between Anaphase-Promoting Complex–Cdh1 and Cyclin A-Cdk2 during Cell Cycle Progression. *Mol Cell Biol* **21**, 3692 (2001).

18. Brandeis, M. & Hunt1, T. The proteolysis of mitotic cyclins in mammalian cells persists from the end of mitosis until the onset of S phase. *EMBO J* **15**, 5280–5289 (1996).

19. Ferrell, J. E. & Ha, S. H. Ultrasensitivity part I: Michaelian responses and zero-order ultrasensitivity. *Trends Biochem Sci* **39**, 496–503 (2014).

20. Ermentrout, B. Simulating, Analyzing, and Animating Dynamical Systems. *Simulating, Analyzing, and Animating Dynamical Systems* (2002) doi:10.1137/1.9780898718195.

21. R Core Team. R: A Language and Environment for Statistical Computing. Preprint at https://www.R-project.org/ (2024).
